# Metagenome-guided substrate selection enriches *Terriglobus*, reveals co-occurring taxa, and enables isolation of a novel species

**DOI:** 10.64898/2026.05.14.725246

**Authors:** Dawn Chiniquy, Spencer Diamond, Hans K. Carlson, Alexey Kazakov, Devin Coleman-Derr, Trent R. Northen, Jillian F. Banfield, Adam M. Deutschbauer

## Abstract

Microbes that remain uncultivated occupy nearly every ecosystem on the planet; this is particularly true in soils, where despite their prevalence, the roles of rarely cultivated microbes in driving biogeochemical cycles and ecosystem function remain poorly explored. We combine metagenome-informed substrate selection with enrichment sub-communities to generate reduced-complexity communities that preserve co-occurrence and expand experimental access to underrepresented soil lineages without requiring prior isolation of each member. Carbohydrate-active enzyme (CAZyme) profiles from soil-derived genomes were used to select carbon compounds predicted to enrich difficult to culture taxa, including members of the phylum Acidobacteriota. Based on 16S rRNA amplicon sequencing, we reproducibly enriched *Terriglobus* (Acidobacteriota) on multiple metagenome-guided substrates. Select communities with consistent presence and varying abundance of *Terriglobus* were passaged in a longitudinal design to generate 89 metagenomes; genus-level profiling revealed that community composition varied between biological replicates but remained consistent within replicates over time, providing diverse Acidobacteriota-containing configurations for downstream analysis. Association network inference identified a core set of co-occurring taxa that positively tracked with *Terriglobus* across the longitudinal series. In parallel, the substrate-guided approach led to isolation of a novel *Terriglobus* species, the first cultured representative of its GTDB species cluster. Together, these results establish a generalizable strategy for generating communities enriched with rarely cultivated taxa, yielding tractable systems for studying microbial interactions and community assembly in soil.

## Introduction

The Earth’s biosphere contains an estimated 1 trillion microbial species [1], which occupy even the most extreme environments; this diversity represents an enormous genomic and metabolic potential that remains mostly untapped, as fewer than 0.1% of known microbial species have been isolated [2]. Advances in sequencing technology have significantly expanded our knowledge of microbial diversity, including the discovery of new phyla, and reshaped our understanding of the tree of life [3]. The remarkable diversity and niche specificity of the microbial kingdom has produced a highly evolved gene repertoire that has been harnessed in human health [4], bioremediation [5,6], plant productivity [7], crop disease resistance [8], industrial biotechnology [9,10], and drug discovery [11,12]. Although cultivation-independent approaches continue to advance our understanding of microbial diversity and function [3], the isolation and cultivation of individual microorganisms, or simplified communities that contain these microorganisms, remains important for experimentally probing microbial ecology [13,14]. Substantial biases in microbial cultivability persist due to factors such as obligate symbioses [15,16], incomplete knowledge of nutritional requirements [17], and long generation times [18].

Soil microbiomes are among the most complex of these ecosystems and are essential to life on Earth, serving as primary drivers of key biogeochemical cycles [19]. Yet our mechanistic understanding of how soil microorganisms interact with one another and with their environment remains limited, presenting a major challenge for large-scale modeling [20]. One strategy to reduce system complexity while preserving essential diversity and ecological relevance is the use of defined microbial consortia [21]. These isolate-based communities have proven particularly valuable in host-associated microbiomes, where they have enabled causal tests of host-relevant traits and microbiome assembly [22,23]. However, a primary drawback of using assembled microbial consortia is that all members must first be isolated. In practice, consortia are often composed of microbes that are not known to interact or may not be isolated from the same environment, possibly limiting ecological relevance [24]. To address these shortcomings, enrichment sub-communities–simplified microbial communities derived from more complex ones under selective pressure–provide a path to study microorganisms that naturally co-occur and interact within a complex system [25–28]. Stable sub-communities have provided insights into human gut microbial dynamics [29,30], microbial element cycling [31–35], and phototroph-associated core microbiomes [36], but have been applied more sparingly to soils [19,37]. Previous efforts to cultivate or enrich soil phyla that are underrepresented in culture collections have largely relied on mimicking oligotrophic, soil-like conditions such as acidic pH, long incubations, and polymeric substrates or dilution-to-extinction, which has enabled isolation of Acidobacteriota and improved recovery and enrichment of Acidobacteriota, Verrucomicrobiota, Chloroflexota, and Gemmatimonadota in reduced-complexity soil communities [37–41].

The goal of this study was to build on recent soil-focused enrichment efforts [28,42–46] by developing stable, reduced-complexity *in vitro* soil sub-communities that included rarely cultivated taxa. To achieve this, we employed a metagenome-guided strategy **[Fig. 1]** to select carbon substrates predicted to enrich members of Acidobacteriota, a phylum abundant and widespread in soils whose members remain comparatively underrepresented among cultured isolates despite prior successes using low-nutrient and polymer-amended media [47,48]. Predictions were based on unique and differentially represented microbial carbohydrate-active enzyme (CAZymes) inventories derived from grassland soil metagenome-assembled genomes (MAGs) [49]. Using the same field soil that informed the metagenomic predictions as inoculum, we initiated enrichment cultures with 20 carbohydrate substrates selected from CAZyme-based predictions, including nine predicted to enrich Acidobacteriota. We identified metagenome-guided substrates that enriched *Terriglobus* (Acidobacteriota), prioritized glucuronoxylan for reproducible enrichment, and used longitudinal genome-resolved metagenomics to track community stabilization and infer co-occurring taxa. This workflow linked metagenome-informed substrate selection to the enrichment and isolation of a novel *Terriglobus* species and provides a framework for studying interactions involving rarely cultivated soil microbes.

**Figure 1.**
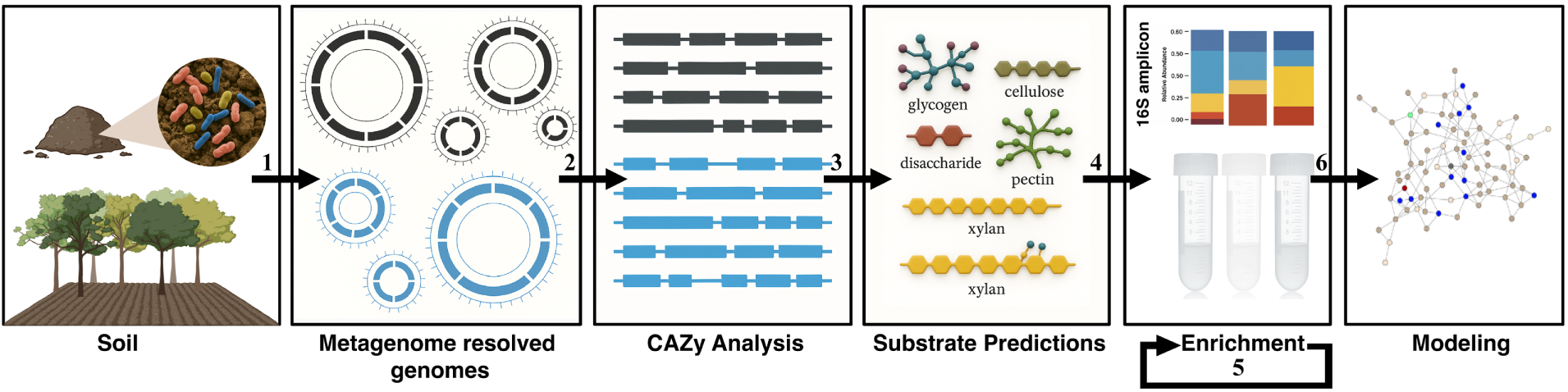
Experimental Overview. The central experimental workflow consisted of the following steps: **1)** Analysis of the genomic content of MAGs recovered from field soil surveys collected at the Angelo Coast Range Reserve; **2)** Global survey of CAZymes in the MAGs associated with recalcitrant taxa from this dataset; **3)** Predictions of substrate preferences for difficult to culture phyla based on CAZy-enzyme content; **4)** Enrichment experiments using select compounds linked to unique CAZy gene activities using the original soil as inoculum source; **5)** Propagation of and optimization of successful enrichment samples containing a consistent presence of difficult to culture taxa; **6**) Longitudinal genome-resolved analysis, resulting in network analysis and modeling of a core microbiome.

## Materials and Methods

### Identification of CAZy associated with rarely cultivated soil lineages from soil metagenome data

Non-Archaeal MAGs with an estimated completeness ≥ 70% and contamination ≤ 5% using CheckM v1.2.1 [50] were retrieved (n = 558 bacterial MAGs) from the dataset of Diamond, et al. 2019 [49]. Proteins in MAGs were predicted with Prodigal v2.6.3 [51] and CAZyme were annotated using the dbCAN v6 HMM database [52]. Statistical analysis of CAZy associations was performed in R v4.3.1. CAZy classes present in at least 2 MAGS were zero-imputed and normalized using centered log ratio (clr). PCA and UMAP were performed directly on the normalized matrix using the prcomp function in R and the umap function from the umap package in R, respectively. Permutational analysis of variance (PERMANOVA) was conducted using the adonis2 function of the vegan package in R. The association between the euclidean distance between genomes in the clr normalized CAZy count matrix and if a genome was a member of one of four soil phyla largely recalcitrant to isolation (Acidobacteriota, Verrucomicrobiota, Chloroflexota, and Gemmatimonadota) was estimated using 9,999 permutations. Filtering thresholds, within-class normalization, and generalized linear model procedures with FDR correction are provided in the Supplementary Methods.

### Field sample collection and processing

Soil was collected in February 2020 from the south meadow field site at the Angelo Coast Range Reserve in Branscomb, California (39° 44′ 21.4′′ N 123° 37′ 51.0′′ W). Soil was collected using a manual coring device with a 1.5 in × 7 in cylindrical polycarbonate insert to remove 10 cm of soil at a time from an individual sampling bore. Soil from 0–10 cm of bore was placed in sterile bags on ice, then transported back to the lab and stored at 4 °C until further processing. The collected soil was passed through a #10 2 mm sieve to remove large particles and plant material.

### Selection of compounds for enrichment experiments

In total 246 unique enzyme classes were analyzed for the following CAZy superclasses (Aux Activity: AA, Carbohydrate Esterase: CE, Glycosyl Hydrolase: GH, and Polysaccharide Lyase: PL). Glycosyl transferases (GT) were omitted from the analysis as their activity is not specific to carbohydrate bond cleavage.

### Enrichment of microbial taxa with carbon compounds

For the targeted enrichment experiments, each compound was dissolved in water and filter-sterilized at 2X concentration. The Freshwater Universal Media (FUM) (Table S10), minimal media containing buffer, salt, minerals, and vitamins was prepared at 4X, adjusting the pH to 5.5 and filter sterilizing. For the untargeted enrichment experiments, the carbon plate was also dissolved in filter-sterilized water, and the phenolic compound plate was dissolved in DMSO or water depending on the compound and as previously described [53–55] (**Table S11**). An 100X amino acid mixture was prepared to further support the growth of microbes on single carbon sources. Initial enrichment cultures were established in 2 ml 96-well deep well blocks and covered with an AeraSeal gas permeable film (Millipore Sigma), with a 400 μl working volume. Once inoculated, cultures were grown in the dark at 30°C with 150 rpm shaking for 10 d. For time series experiments, each transfer to new media was a 1:10 subculture, transferring 40 μl of enrichment culture to 360 μl of new growth media. With each transfer and sample collection for DNA isolation, 20% glycerol stocks were made of the cultures to enable retrieval of individual samples that had the highest abundance of difficult to culture microbes once profiled by 16S rRNA sequencing. Substrate dissolution recipes, individual well composition, soil extract preparation [56,57], the salt-stressed variant, and DNA pellet storage are provided in the Supplementary Methods.

### DNA extraction, Amplification, and Illumina Sequencing of 16S Sequences

Genomic DNA was extracted from samples using the 96-well Qiagen DNeasy Blood & Tissue kit (catalog no. 69516), following the manufacturer’s protocol for Gram-positive bacteria. The V4/V5 region of the bacterial 16S rRNA gene was PCR amplified from extracted DNA using the 515F/926R primers with in-line dual Illumina indexes [58]. The resulting libraries were quantified using the Qubit dsDNA BR Assay kit (Invitrogen, catalog no. Q32853), pooled, cleaned using the Zymo Research Clean and Concentrator kit (Zymo Research, catalog no. D4014), and sequenced on an Illumina MiSeq (PE 2x300). For 16S read processing, paired-end reads were merged using PEAR [59], demultiplexing using inline indexes, and reads with more than one expected error was detected and discarded, as well as rare sequences (fewer than 4 reads in any sample) using USEARCH [60]. Chimera removal and error correction were performed with UNOISE3 [61]. Taxonomy was assigned using SINTAX [62] with the RDP training set v18 [63].

### Assembly, binning, dereplication, and genome quality

Raw metagenomes from 89 samples were assembled with IDBA-UD [64] and SPAdes [65]. Contigs were binned using four independent binners, and bins were consolidated with DASTool [66]. For each bin, taxonomy was assigned with GTDB-Tk [67], and completeness and contamination were estimated with CheckM [50]. Species-level dereplication used dRep [68] at ANI ≥95% to select a single species-representative bin per cluster, maximizing completeness and contiguity while minimizing contamination. We predicted coding sequences with Prodigal [51], rRNA and tRNA genes with standard workflows, and recorded codon usage with Codetta [69]. The final set comprised 106 species-representative genomes with high quality (mean completeness 95.9%, mean contamination 1.15%; **Table S7**).

### Read mapping and abundance profiling

Quality-controlled reads were competitively mapped to the 106-genome representative set to obtain per-species coverage and relative abundance per sample. Mean mapping rates were ∼80% across the dataset, consistent with expectations for reduced-complexity enrichment communities. To compare enrichments and derived stocks, we computed alpha diversity (observed species) and constructed ordinations on CLR-transformed relative abundance matrices. *Core microbiome definition and network inference*

We defined prevalence as the fraction of enrichment samples (n=71) in which a species was detected. The core microbiome was the set of species present in ≥95% of enrichments. We also reported species present in 100% of enrichments and compared core membership between biological replicates of enrichment communities over time. We filtered to species present in ≥10 enrichments (n=71) to reduce sparsity. We inferred two complementary networks: SparCC correlations from relative abundance data with permutation to assess edge significance [70] and SPIEC-EASI conditional-dependence graph [71] using neighborhood selection with StARS for model selection [72]. We retained positive edges and defined a consensus network by requiring presence in both SparCC and SPIEC-EASI [71,72]. We summarized node-level statistics (degree, betweenness, eigenvector centrality) and global properties using igraph [73]. To identify community structure, we applied the weighted Leiden algorithm [74] with modularity optimization across resolution values ranging from 0.1 to 1.5 (step = 0.05), selecting the resolution that maximized modularity (resolution = 0.75, modularity = 0.67). To assess clustering stability, we performed 100 independent Leiden iterations at the optimal resolution using different random seeds; 79% of runs recovered seven communities with consistent modularity scores (mean = 0.67 ± 0.004). Leiden clustering-stability details are provided in the Supplementary Methods.

### CAZyme annotation of enrichment MAGs

To assess hemicellulose-degrading capacity among direct network neighbors and core microbiome members, we predicted proteins with Prodigal v2.6.3 and searched against Pfam HMMs for GH43 (PF04616) and GH62 (PF03664) using HMMER v3.4 with an E-value threshold of 1 × 10⁻⁵.

### Isolation of Terriglobus sp. from enrichment cultures

Enrichment cultures were re-grown from glycerol stocks of the communities stored in a - 80 °C freezer by inoculating 3 mls of FUM minimal media pH 5.5 with glucuronoxylan (50 mg/40 ml) with growth for 10 d, serially diluted to 10-4 and spread on agar plates composed their respective growth media. Individual colonies were collected and re-streaked on the same media. Isolation of the *Terriglobus* sp. DMC71 required over 6 rounds of re-streaking to get a pure culture. Iterative 16S profiling and re-streaking of mixed cultures between rounds are provided in the Supplementary Methods.

### SEM imaging of Terriglobus sp. DMC71

Scanning electron microscopy preparations (Electron Microscopy Laboratory, University of California, Berkeley) were: *Terriglobus* sp. DMC71 isolate was grown in 0.1X R2A for 7 d (shaking at 150 rpm at 30°C, then gently pelleted, and resuspended in a fixative solution (2% glutaraldehyde in 0.1 M sodium cacodylate buffer, pH 7.2) and was placed at 4°C rotating for 3 hr then stored at 4°C in dark (standing) overnight. Samples were then transferred onto 0.1% (w/v) poly-L-lysine–coated coverslips and subjected to critical point drying (Tousimis, Mayland, USA). SEM imaging was performed using a Zeiss Crossbeam 550 (Carl Zeiss Microsystems GmbH, Oberkochen, Germany). Post-fixation, washing, dehydration, and sputter-coating protocols are provided in the Supplementary Methods.

## Results

### Global analysis of CAZymes associated with rarely cultivated soil lineages

Several of the most abundant phylum-level taxa identified in global soil surveys [46,75] are underrepresented among cultivated isolates [41], limiting species-level investigation and modeling of soil microbiome dynamics. To quantify this cultivation gap, we examined the prevalence of Latinized species names in the Genome Taxonomy Database (GTDB-R226) across ten globally abundant soil phyla, using Latinized nomenclature as a proxy for species-level isolation; Acidobacteriota, Verrucomicrobiota, Chloroflexota, Gemmatimonadota, and Planctomycetota were significantly under-represented among named isolates and referred to here as the recalcitrant group (LOR < 0, FDR ≤ 0.001; **Fig. S1**). We then re-analyzed MAGs from Angelo Coast Range Reserve grassland soils [49]; across 558 bacterial MAGs with sufficient quality (Completeness ≥ 70% and Contamination ≤ 5%; **Tables S1-S3**), 240 unique CAZyme classes were detected. CAZyme inventories of MAGs were significantly associated with phylum (*p*-value = 3e^-4^, R = 0.17, n = 9,999 iterations; PERMANOVA). Principal component analysis (PCA) exhibited a horseshoe pattern indicative of a non-linear relationship **[Fig. S2A and B]**, whereas Uniform manifold approximation (UMAP) resolved phylum-level distinctions **[Fig. 2A]** and separated MAGs by over- and under-representation in culture collections **[Fig. S3A and B]**, indicating that CAZyme content varies with phylum and cultivability.

**Figure 2.**
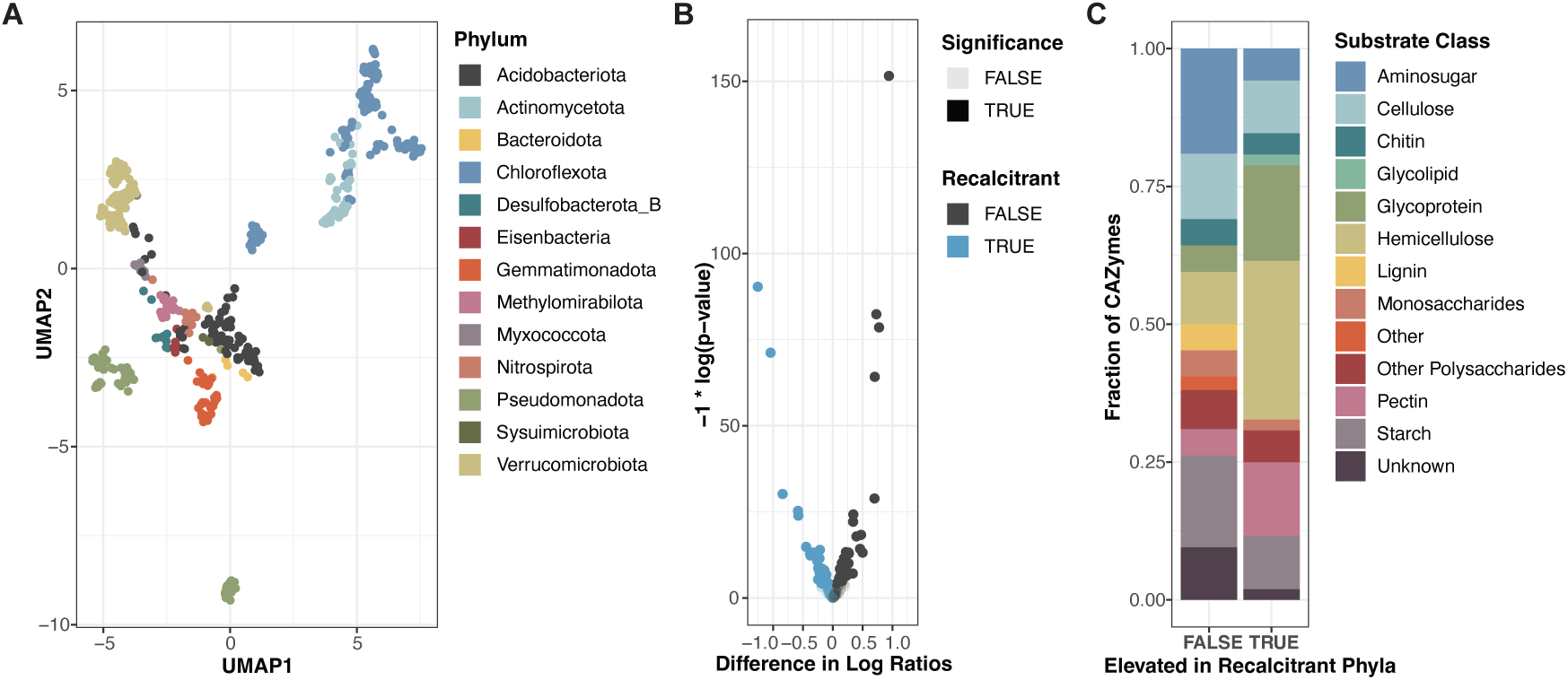
Comparative CAZyme profiling across MAGs reveals differential enzyme and substrate-class representation between phyla over- and under-represented in culture collections. **(A)** UMAP projection of centered log-ratio (CLR)-normalized CAZyme counts across 558 high-quality bacterial MAGs (>70% completeness, <5% contamination) from a previously published Angelo Coast Range Reserve grassland soil metagenome dataset (Diamond et al. 2019). Each point represents a single MAG and is colored by phylum based on GTDB-R226 taxonomy. **(B)** Volcano plot of CAZyme classes differentially abundant between MAGs from recalcitrant (under-represented in named isolates; **Fig. S1**) and non-recalcitrant phyla. The x-axis shows the CLR effect size (recalcitrant minus non-recalcitrant) and the y-axis shows - log10(FDR). Each point represents one CAZyme class; significance was defined as ANOVA FDR ≤ 0.05 and contrast p ≤ 0.05. A total of 80 CAZyme classes differed significantly, of which 33 were enriched in recalcitrant phyla. **(C)** Predicted substrate classes targeted by significantly differentially abundant CAZyme classes, separated by whether the class was enriched in recalcitrant or non-recalcitrant phyla. Hemicellulose- and glycoprotein-targeting CAZymes dominated the recalcitrant-enriched subset.

We identified 80 CAZyme classes that differed significantly between phyla that are underrepresented versus well represented in culture collections; 33 CAZyme classes were enriched in the underrepresented (recalcitrant) group **[Fig. 2B, Table S4]**. Among the recalcitrant-enriched subset, predicted target substrates were dominated by hemicelluloses and glycoproteins, whereas CAZymes enriched in the well-represented group more often targeted starch, pectin, and amino-sugar polymers **[Fig. 2C]**. We also identified 62 CAZyme classes that were phylum-specific (present only within a single phylum in this dataset) **[Table S5]**, with the largest numbers in Acidobacteriota (n = 14 classes), Bacteroidota (n = 16 classes), and Chloroflexota (n = 14 classes). Xylan deconstruction emerged as a recurring functional signature of recalcitrant phyla, with evidence for both broadly enriched and lineage-specific CAZyme families; GH3, which includes β-glucosidase/β-xylosidase activity on glucuronoxylan, and GH51, an α-L-arabinofuranosidase involved in arabinoxylan debranching, were enriched in recalcitrant phyla **[Table S4]**, while the phylum-specific subfamilies GH30_2/7, associated with endo-β-1,4-xylanase/glucuronoxylanase activity, and GH43_11, associated with β-xylosidase and α-L-arabinofuranosidase activity, occurred only in Chloroflexota and Acidobacteriota, respectively [**Table S5**]. We then matched these CAZymes to commercially available substrates by manufacturer-reported primary linkages **[Table S6]**; nine compounds were predicted to enrich Acidobacteriota, including xylans, mannans, arabinogalactan, chondroitin, and rhamnogalacturonan-I [**Fig. S4A**].

### Substrate-guided enrichment of rarely cultivated soil lineages

We established enrichment cultures using 20 carbon compounds linked to CAZymes enriched in rarely cultivated lineages [**Table S6**], including nine predicted to enrich Acidobacteriota. Using field soil from the metagenomic study site as inoculum, cultures were grown in defined media at acidic pH, with and without a high salinity treatment [37]. Enrichment cultures were harvested once visually turbid (10 d) and their community composition was analyzed via 16S rRNA amplicon sequencing. Five bacterial phyla were detected across samples, with fewer than 29 observed ASVs per sample [**Fig. 3A**]. Among rarely cultivated phyla, Acidobacteriota was observed in cultures containing xylans, mannans, and galactomannan (PERMANOVA R² = 0.26, p = 0.001), validating four of nine metagenome-based predictions [**Fig. 3B**]. Alginic acid, glycogen, and rhamnogalacturonan-I were predicted to enrich Acidobacteriota and Bacteroidota, but enrichments with these compounds did not yield Acidobacteriota reads. No reads were recovered for Verrucomicrobiota, Gemmatimonadota, or Chloroflexota under any condition, and high-salt cultures greatly favored Actinomycetota growth **[Fig. S4B and C]**.

**Figure 3.**
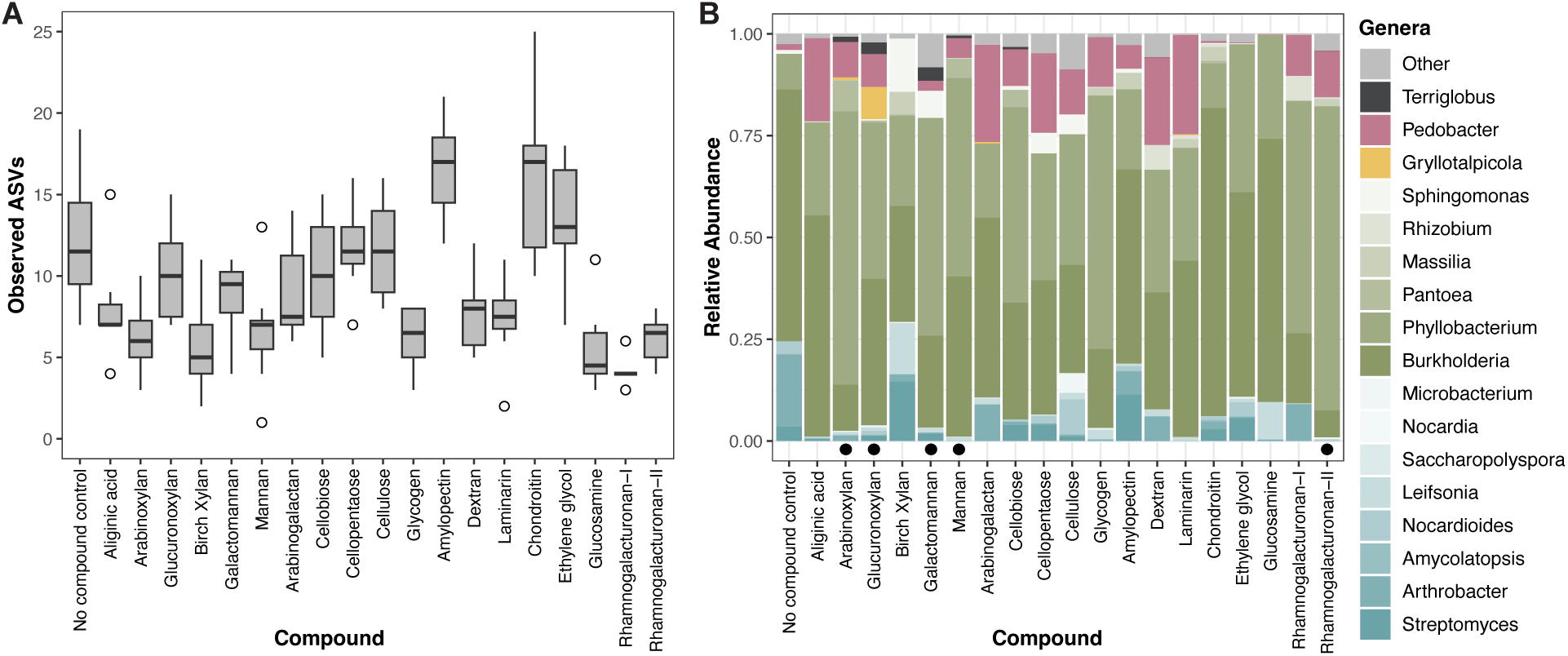
Enrichment of microbial communities using soil inoculum from original study and predicted carbon substrates from CAZy metagenome catalog. **(A)** Observed ASVs per compound based on 16S rRNA amplicon sequencing. Observed ASVs plotted across each of the treatment groups (n=20), demonstrating a significant impact of treatment on alpha diversity (ANOVA, *p*<2e-16). Boxplots show medians, interquartile ranges, and whiskers extending to 1.5 × IQR. **(B)** Mean phylum-level relative abundance across biological replicates (n=8) for each compound. Black circles indicate compounds for which at least one biological replicate met the Acidobacteriota-positive threshold (≥5 reads per 1,000 total reads). Among predicted Acidobacteriota-enriching substrates, xylans, mannans, and galactomannan supported Acidobacteriota enrichment.

To benchmark this method against an untargeted enrichment approach, we screened 192 compounds including monosaccharides, polysaccharides, amino acids, nucleic acids, and aromatic acids, distributed across a carbon plate and a phenolic compound plate [32,76]. For each compound, we performed 4 replicates at neutral pH with otherwise identical media conditions. Most substrates supported commonly cultured phyla (Actinomycetota, Bacteroidota, and Pseudomonadota) [**Fig. 4A and 4B, Fig. S5**]. In contrast to the targeted screen focused on Acidobacteriota, the broader compound screen yielded two additional recalcitrant phyla: Gemmatimonadota and Planctomycetota **[Fig. S5**], with additional substrates identified that enriched for Acidobacteriota (3-hydroxybutyric acid, mandelic acid, and vanillyl alcohol). The fraction of Acidobacteriota-positive samples (≥5/1000 reads) was similar between screens (targeted: 5.9%; untargeted: 4.0%; χ²=0.52, *p*=0.47), but the targeted approach yielded ∼10-fold higher number of average Acidobacteriota reads per positive sample (47.9 vs 6.0) [**Fig. 4A and 4B]**.

**Figure 4.**
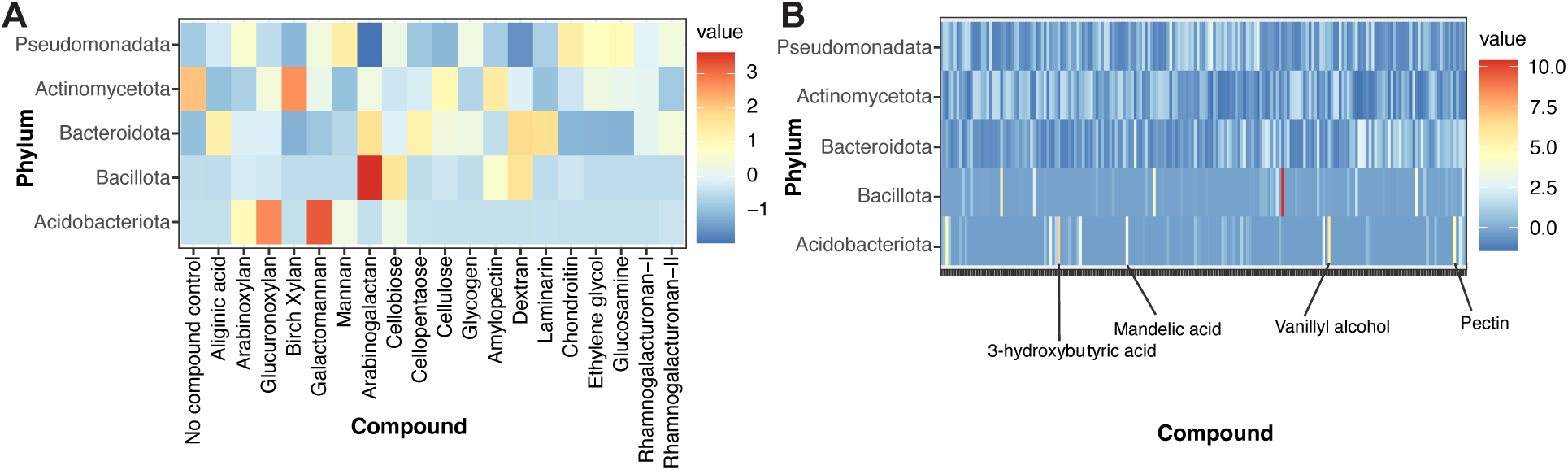
Phylum-level abundance distributions across CAZy-targeted and untargeted screening approaches. **(A)** Mean relative abundance of total reads belonging to each phylum (y-axis) observed across each compound (x-axis) based on 16S rRNA amplicon sequencing. Data represents average of total read counts across a total of 8 biological replicates per compound. **(B)** Mean relative abundance of total reads belonging to each phylum (y-axis) observed across each of 192 compounds (x-axis) based on 16S rRNA amplicon sequencing. The broader screen recovered additional Acidobacteriota-positive substrates, including 3-hydroxybutyric acid, mandelic acid, and vanillyl alcohol.

### Optimization and selective propagation of Acidobacteriota enrichment cultures

To optimize Acidobacteriota growth, we retested six substrates that yielded Acidobacteriota in the initial screen plus their admixture (arabinoxylan, glucuronoxylan, mannan, galactomannan, cellobiose, and cellopentaose), under modified conditions (increased replication, additional timepoints, static versus shaking) [47,77]. Acidobacteriota reads were largely absent in static conditions but recovered under shaking despite similar observed ASV ranges **[Fig. S6A-B]**, indicating sensitivity to oxygen transfer. Glucuronoxylan, mannan, galactomannan, and cellobiose produced the highest Acidobacteriota abundance at 14 d, and the combination of substrates did not improve Acidobacteriota enrichment **[Fig. S6C**]. In parallel, glucuronoxylan and galactomannan (locust bean gum (LBG)) enrichments (n = 18) were used to test optimal storage and recapitulation conditions.

To identify a reproducible substrate for longitudinal metagenomics, we ranked Acidobacteriota-positive enrichments (≥5/1000 reads; n=27); galactomannan yielded the two highest relative abundances of Acidobacteriota (50.4%, 29.7%) but was inconsistent across replicates (39/42 negatives) **[Fig. 5A]**, whereas glucuronoxylan was the most reproducible (17/40 positives; 7/8 biological replicates positive). We therefore advanced eight glucuronoxylan-derived communities spanning high and low Acidobacteriota abundance (**Fig S7A-B**). Storage and recapitulation tests on one high abundance community **[Fig. 5B-C]** supported 30% glycerol cryostorage and continued use of FUM medium at pH 5.5 for downstream metagenomics.

**Figure 5.**
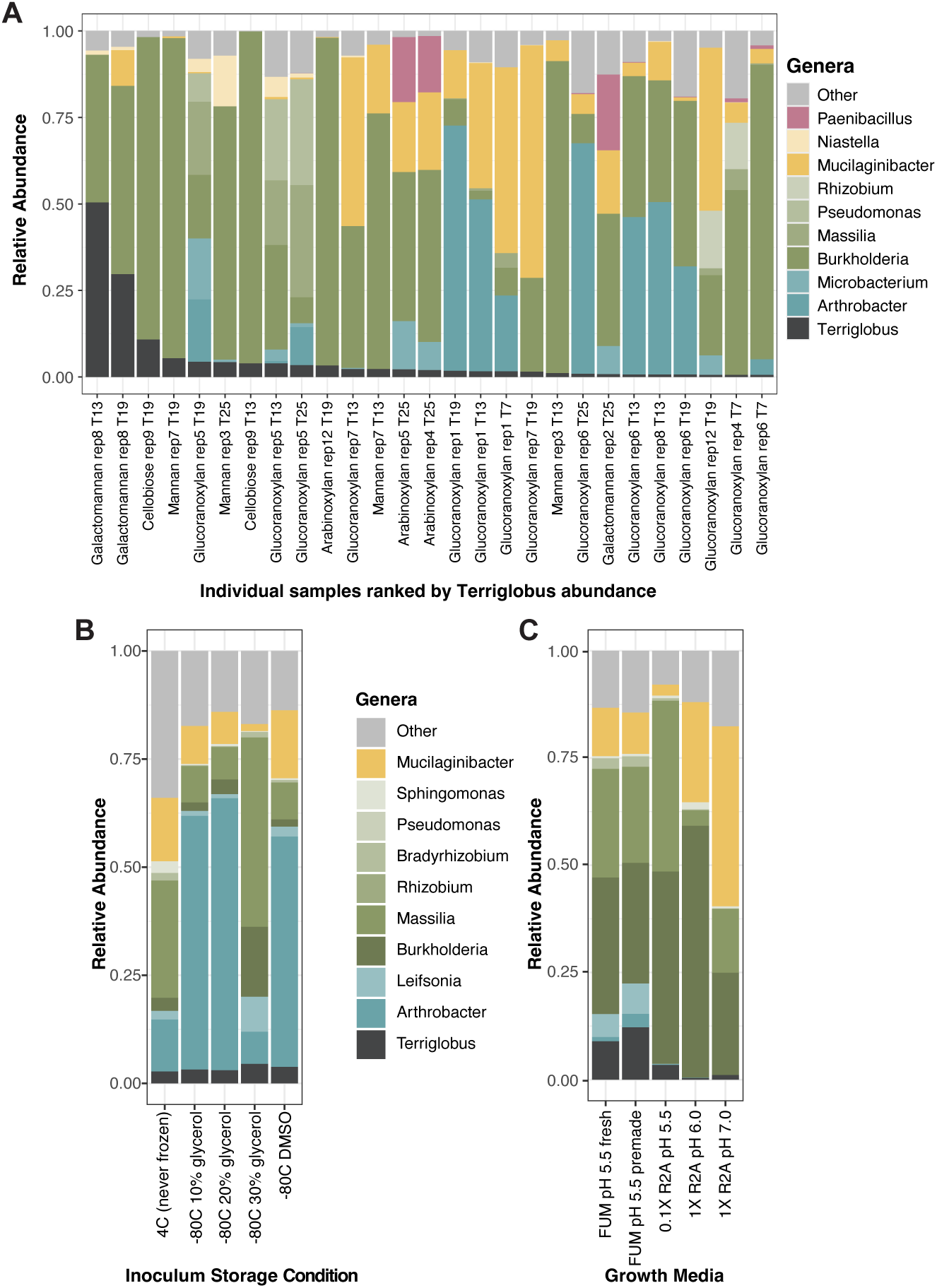
Substrate selection and growth optimization for metagenomic analysis based on 16S rRNA amplicon sequencing. **(A)** Ranked Acidobacteriota-positive enrichments (≥0.5% relative abundance; n = 27) based on 16S rRNA amplicon sequencing. Among the 27 Acidobacteriota-positive enrichments, galactomannan yielded the highest relative abundances, but glucuronoxylan was the most reproducible substrate overall (17/40 positives; 7/8 biological replicates positive) and was therefore selected for longitudinal metagenomics. **(B)** Storage-condition tests for glucuronoxylan-derived communities. Cryostorage at −80 °C in 30% glycerol yielded the highest *Terriglobus* relative abundance. **(C)** Media optimization tests for glucuronoxylan-derived communities. Freshwater Universal Medium (FUM) at pH 5.5 was selected for downstream metagenomics. Values in panels B and C are means across 4 biological replicates.

### Longitudinal metagenome analysis of enrichment cultures containing Acidobacteriota

As compositional variation aids both binning and network inference [27,78–80], we designed a longitudinal series to maximize variation in Acidobacteriota abundance and community composition. First-stage passaging sampled eight glucuronoxylan enrichments (four high and four low Acidobacteriota) at days 7, 14, 21 [**Fig. S7]**, and glycerol stocks from each timepoint served as inoculant for second-stage passaging on the same timeline and media. This yielded 72 metagenomes that, with 18 glycerol-stock test samples, totaled 89 metagenomes (mean depth 14.1 Gb/sample). Acidobacteriota were detected in all enrichment cultures. Genus-level relative abundance profiles varied substantially between biological replicates but remained largely consistent within each replicate across the 21-day time course regardless of which Stage 1 glycerol stock served as inoculum, indicating that assembly was driven primarily by initial replicate identity rather than passage timepoint. The four biological replicates initially selected for high Acidobacteriota abundance maintained elevated *Terriglobus* levels throughout the Stage 2 passaging (asterisks, **Fig. 6A**). Over time, communities converged toward a stable structure, with Acidobacteriota emerging even from inocula where they were initially rare or undetected, consistent with proliferation from the rare biosphere under selection (**Fig. 6B**).

**Figure 6.**
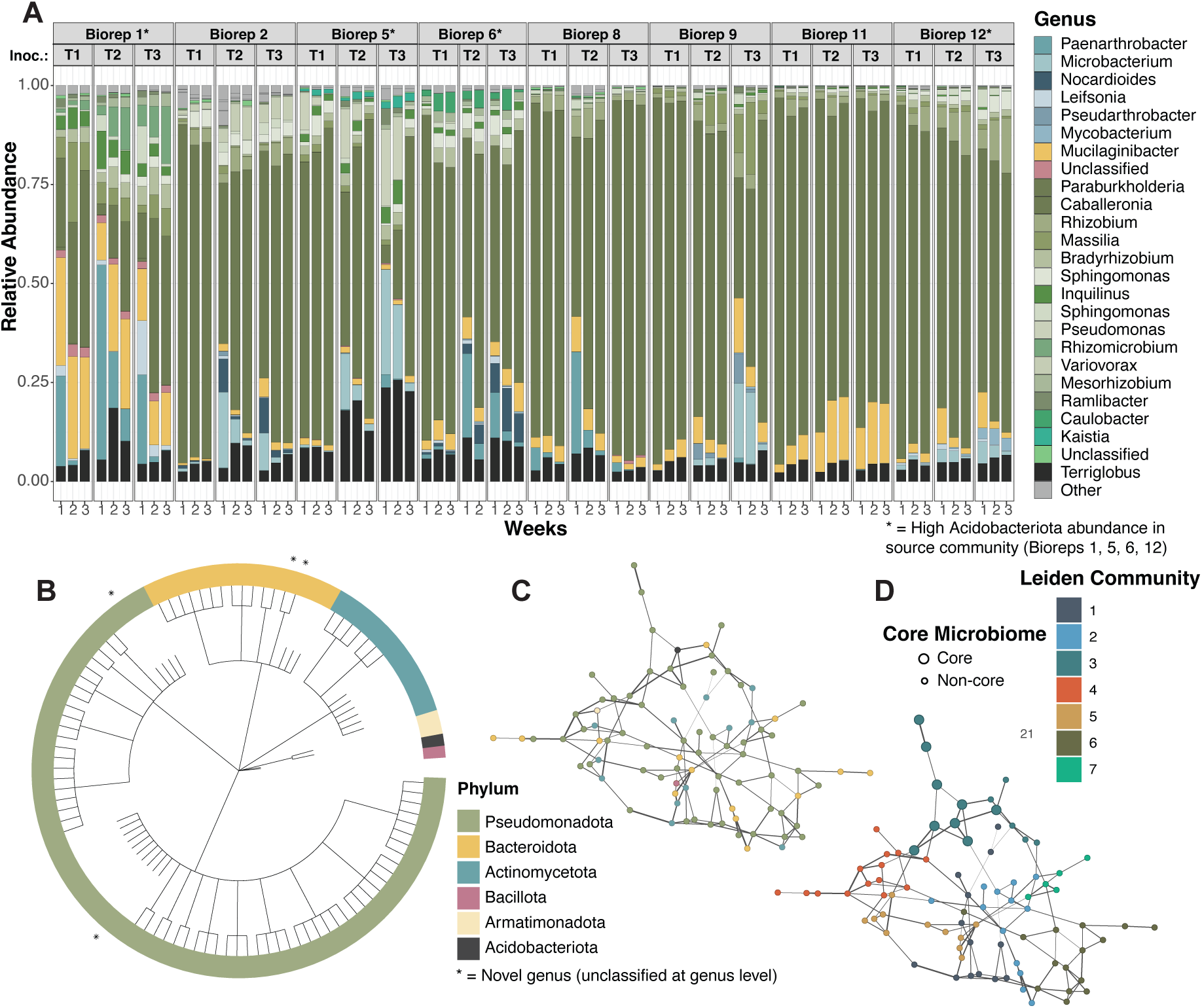
Genome-resolved metagenomics and network analysis of glucuronoxylan-derived communities. First-stage passaging of eight glucuronoxylan-derived communities generated inocula for second-stage passaging and longitudinal shotgun metagenomics. A total of 89 metagenomes were analyzed. **(A)** Genus-level relative abundance profiles of 106 MAGs across 72 second-stage metagenomes derived from eight biological replicates, three Stage 1 inoculum timepoints (T1, T2, T3 weeks), and three Stage 2 sampling timepoints (1, 2, 3 weeks). Asterisks denote the four communities initially selected for high Acidobacteriota abundance based on 16S profiling. Composition varied substantially among biological replicates but remained relatively consistent within each replicate across Stage 2 sampling timepoints and Stage 1 inoculum sources. **(B)** Circular phylogenetic tree of the 106 species-representative MAGs, colored by phylum. Asterisks mark MAGs lacking genus-level classification in GTDB: Chitinophagaceae (n = 2), Burkholderiaceae (n = 1), and Beijerinckiaceae (n = 1). The average genome size was 5.4 ± 1.3 Mbp and the average completeness was 95.9 ± 6.9%. **(C)** Consensus co-occurrence network inferred from SparCC and SPIEC-EASI analyses of the longitudinal glucuronoxylan-selected communities (n = 71 samples), comprising 85 nodes and 136 edges. Node colors indicate phylum-level taxonomy. **(D)** Community structure inferred using the Leiden algorithm at a modularity-optimized resolution of 0.75 (modularity = 0.67), revealing seven communities. Larger nodes denote the nine core microbiome members (present in ≥95% of samples), all of which clustered within a single Leiden community.

### Leveraging metagenomic analysis to uncover the core microbiome in Acidobacteriota-enriched microbial communities

Assembly, binning, and dereplication of the longitudinal series yielded 106 species-representative metagenome assembled genomes (MAGs) (ANI ≥95%; mean completeness 95.9%, mean contamination 1.15%), with novel genera marked on the phylogenetic tree (**Fig. 6B**). A single *Terriglobus* species aggregated 88 contributing MAGS, indicating broad occurrence. Competitive read mapping recruited ∼80% of reads per sample. A core microbiome was defined as species detected in ≥95% of enrichments (n=71). Nine species met this threshold, seven of which including the Acidobacteriota member were present in all enrichments **[Table S8**]. To identify taxa co-occurring with Acidobacteriota, we inferred a species-level consensus association network from positive, sign-consistent SparCC and SPIEC-EASI edges **[Fig. 6C]**. The network comprised 85 nodes and 136 edges, with SparCC and SPIEC-EASI edge weights highly correlated (*p* < 2.2 × 10⁻^16^; **Fig. S9**). Direct neighbors of Acidobacteriota were *Niastella* (Chitinophagaceae), *“*BOG-938*”* (Caulobacteraceae), and *Bradyrhizobium* (Xanthobacteraceae); none resolved to a named species. HMM annotation revealed that *Niastella* encoded 13 GH43 family enzymes (hemicellulose debranching), whereas the two *Bradyrhizobium* neighbors lacked detectable GH43 or GH62 homologs, consistent with distinct functional roles **[Table S9]**. Leiden community detection (modularity-optimized resolution 0.75; modularity 0.67) identified seven communities. Notably, all nine core microbiome members clustered in a single community, indicating that the core taxa form a cohesive network module rather than bridging disparate groups [**Fig. 6D**].

### Isolation and imaging of Terriglobus sp. from enrichment cultures

From glycerol stocks of communities with consistently high abundance of Acidobacteriota, six rounds of streaking yielded a *Terriglobus* sp. from glucuronoxylan enrichments. SEM showed short, ellipsoidal rods (∼1 × 0.6 µm) with a pronounced rugose surface (ridges and furrows) and no visible appendages **[Fig. 7]**. This texture was unique among isolates processed and imaged in the same batch, indicating an intrinsic cell-surface feature rather than a preparation artifact. Morphology and size matched prior descriptions for *Terriglobus* [39], but SEM reports of ridged envelopes are lacking, although thin-section electron micrographs in Eichorst et al. (2007) are consistent with our observations [39]. Overall, the observed morphology is consistent with published descriptions of *Terriglobus* as short, non-motile rods [81–83], while documenting a rugose cell surface in the novel strain.

**Figure 7.**
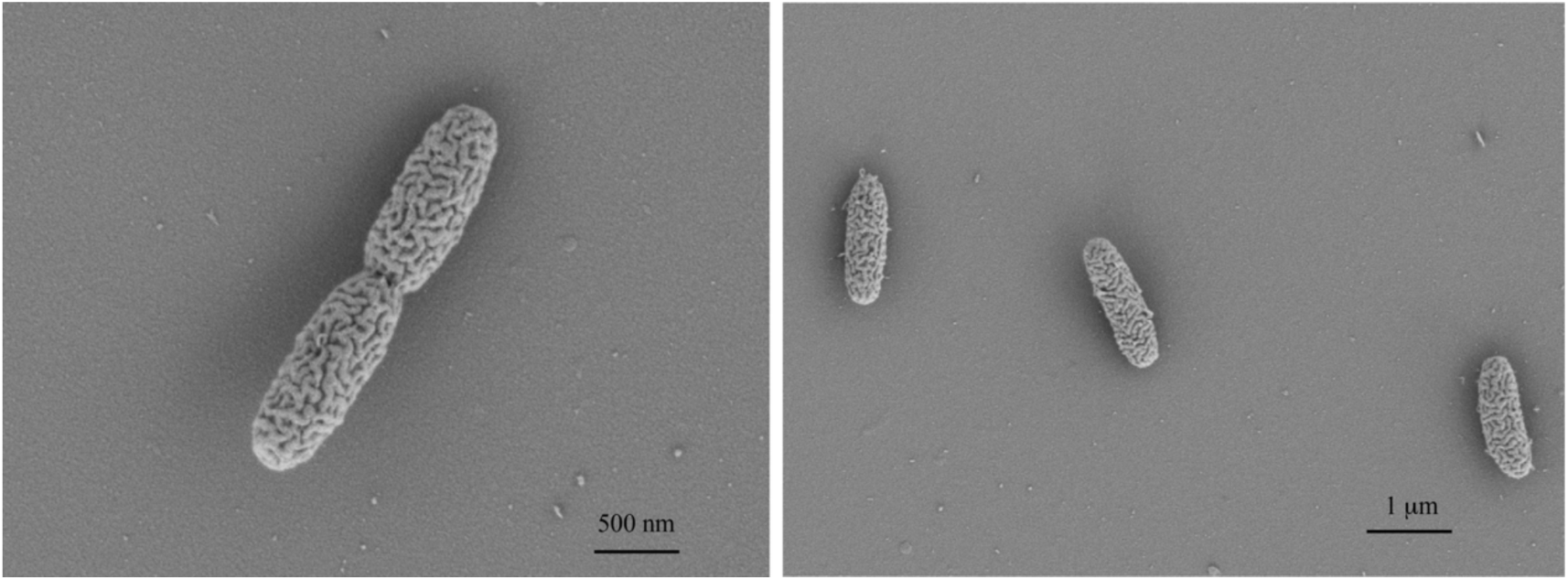
Scanning Electron Microscopy (SEM) imaging of a novel *Terriglobus* sp. isolated from reduced-complexity enrichment communities. *Terriglobus* sp. DMC71 was isolated from a glucuronoxylan-enriched microbial community that showed consistent Acidobacteriota presence by 16S rRNA amplicon sequencing. Cells were grown for 7 d in 0.1× R2A before SEM imaging. DMC71 cells were short, ellipsoidal rods (∼1 × 0.6 µm) with a pronounced rugose surface and no visible appendages. The observed morphology is consistent with published descriptions of *Terriglobus* as short, non-motile rods while documenting a rugose cell surface in strain DMC71.

Hybrid short and long-read sequencing of *Terriglobus* sp. DMC71 produced a closed 6.15 Mb genome (57.9% GC; 100% complete, 0% contaminated by CheckM) encoding 5,035 protein-coding genes, 52 tRNAs, and a single rRNA operon. GTDB-Tk v 2.4.0 [84,85] placed DMC71 in species cluster *Terriglobus_A* sp019510535 (98.5% ANI, 92% alignment fraction), previously represented only by a MAG (SH_60_12) with no cultured representative, making *Terriglobus* sp. DMC71 the first isolate of this species and a candidate type strain. Its closest named relative is *T. albidus* (90.8% ANI; **Fig. S8**), below the 95% species threshold. Consistent with its enrichment on xylan-derived substrates, *Terriglobus* sp. DMC71 possesses a substantial hemicellulose-degrading enzyme repertoire: 4 GH10 endo-xylanases, 12 GH43 family enzymes (including arabinan endo-1,5-α-L-arabinosidases), 3 GH51 α-L-arabinofuranosidases, and 6 GH3 β-glucosidases, consistent with the GH43 content observed in the corresponding enrichment MAG (**Table S8**). Although no flagella were seen by SEM, the genome encodes a complete flagellar biosynthesis apparatus (36 genes) and chemotaxis system, suggesting condition-dependent motility not expressed under the imaging conditions.

## Discussion

Across grassland soil MAGs [49], recalcitrant phyla accounted for most phylum-specific CAZy enrichments, with Acidobacteriota and Chloroflexota contributing the greatest number of recalcitrant-associated CAZy families. The predicted substrates targeted by these repertoires, particularly hemicelluloses and glycoproteins, point to specific resource preferences and subfamily-level specialization, consistent with physiologies adapted to complex polymers that are not typically represented in standard laboratory media [17,27]. This mechanistic context can help explain persistent cultivation gaps and slow growth when key nutrients, cofactors, or native polymer architectures are missing [17,18,47]. In line with these expectations, our CAZyme-guided enrichments reproducibly increased *Terriglobus* (Acidobacteriota) abundance on glucuronoxylan and galactomannan, consistent with the idea that presenting relevant polymers can surface otherwise latent catabolic capacity. This is consistent with the structural complexity of substituted xylans, whose degradation requires coordinated backbone cleavage, side-chain removal, uptake, and intracellular metabolism by enzymes such as endoxylanases, β-xylosidases, α-L-arabinofuranosidases, glucuronidases, and esterases [86,87]. *Terriglobus* sp. DMC71 encodes predicted GH10, GH43, GH51, and GH3 enzymes, linking the source-MAG signal to the isolate-level repertoire. Glucuronoxylan may also have shaped community assembly by favoring xylan specialists over fast-growing generalists and by selecting partners that process released arabino-/xylo-oligosaccharides, consume monomers or organic-acid byproducts, or relieve cofactor limitations [88,89].

However, not all predicted substrates yielded Acidobacteriota growth, despite the presence of putatively matching CAZy annotations in source MAGs. These discrepancies may arise from polymer chemistry, regulatory context, and annotation uncertainty; side-chain composition, degree of polymerization, and linkage heterogeneity can require specific debranching enzymes and transporters whose expression may be conditional, while CAZy family-level annotations may not fully resolve substrate specificity, catalytic activity, or enzyme functionality under the tested conditions [26,27]. Abiotic parameters further shaped outcomes: high salt shifted communities toward Actinomycetota while acidic pH favored Acidobacteriota, in line with prior observations [37,38,47]. Variation among replicate enrichments likely further contributed to these outcomes. Between-replicate variance in these low-nutrient, reduced-diversity enrichments is consistent with stochastic assembly, priority effects, and ecological drift, with most of the divergence in our dataset likely arising during the initial bottleneck in each biological replicate, as genus-level composition remained largely consistent within each replicate over time despite substantial differences between replicates [90,91]. Together, these observations emphasize that media composition and physical regime (oxygen transfer, pH, ionic strength, and passaging history) can override substrate-level predictions and therefore must be co-optimized when the goal is to increase a focal lineage [26,44,56,92].

Across the longitudinal series, glucuronoxylan-fed biological replicates diverged in overall community composition yet repeatedly recovered the same core taxa and *Terriglobus*-associated neighbors, including Niastella, “BOG-938,” and Bradyrhizobium. This pattern suggests that the core microbiome and *Terriglobus*-centered associations were not confined to a single narrowly defined community state but emerged across multiple related *Terriglobus*-containing configurations. The clustering of these highly prevalent taxa within a single Leiden community further indicates that they occupy one cohesive region of the inferred association structure [79,80]. Together, these patterns nominate a recurring co-occurrence context for *Terriglobus* during glucuronoxylan-driven assembly. Whether these repeated associations reflect shared abiotic preferences, trophic coupling, or both cannot be resolved from co-occurrence data alone.

This co-occurrence structure is compatible with a distributed degrader–consumer–facilitator framework rather than a model in which a single taxon performs all steps of polymer turnover. The *Terriglobus* genome encoded substantial GH43 content, consistent with direct participation in hemicellulose deconstruction, while the *Niastella* neighbor encoded 13 GH43 homologs and may provide complementary debranching or oligosaccharide-processing activities, suggesting that *Terriglobus* and *Niastella* may occupy partially overlapping but distinct roles during glucuronoxylan turnover, with each contributing to the release or processing of arabino- /xylo-oligosaccharides from substituted xylans [86]. Co-occurring Alphaproteobacteria (e.g., *Bradyrhizobium* and the Caulobacteraceae lineage) are plausible consumers of monomers and low-molecular-weight byproducts generated during polymer turnover, potentially influencing community stability through resource partitioning and byproduct removal rather than through direct polymer deconstruction [87,93–96]. Among the nine core microbiome members, *Terriglobus* was the only taxon encoding substantial GH43 content (12 enzymes; **Table S8**), consistent with its role as the primary hemicellulose degrader in the consortium. The association of CAZyme-poor taxa with *Terriglobus* could reflect: (i) vitamin/cofactor exchange that relieves auxotrophies [15,16,97], (ii) removal or detoxification of inhibitory intermediates generated during polymer turnover [88], and/or (iii) spatial or metabolic niche partitioning that reduces direct competition and stabilizes coexistence [94], in addition to (or instead of) direct polymer deconstruction. Overall, these associations support a model of distributed, modular organization rather than dominance by a single hub taxon, consistent with patterns reported in other tractable enrichment systems [25–28,89].

The enrichment pipeline also supported classical isolation, yielding a *Terriglobus* sp. that represents the first cultured member of GTDB species cluster *Terriglobus_A* sp019510535—a lineage previously known only from a single MAG. This isolate expands the cultured diversity of *Terriglobus* and provides a candidate type strain for formal species description. Repeated streaking from glucuronoxylan communities yielded a *Terriglobus* sp. SEM revealed short, ellipsoidal rods with a distinctive rugose surface ornamentation. Morphology and size are consistent with the genus, and the rugose envelope is, to our knowledge, the first such surface morphology documented by SEM for the genus [39]. The ability to move from enrichment and network-informed community context to pure culture closes the loop from genome-informed prediction to a tractable culture for downstream functional assays.

Although Acidobacteriota were the focal lineage of this study, the broader 192-substrate screen revealed informative patterns for other phyla. Gemmatimonadota were strongly enriched on dextran, xanthan, and several lactones suggesting specialization on extracellular polysaccharides and their breakdown products [98,99]. Conversely, expected growth by Gemmatimonadota or Verrucomicrobiota to wheat arabinoxylan were lacking under our acidic regime, underscoring how media conditions and polymer heterogeneity complicate predictions drawn from genome inventories alone. More generally, raising the relative abundance of a specific lineage may require the alignment of substrate, abiotic regime, and ecological context, central themes in enrichment design and microbiome engineering [26,27,92].

In sum, metagenome-informed substrate selection grounded in field-derived CAZyme inventories reproducibly enriched the recalcitrant Acidobacteriota lineage *Terriglobus*. Co-optimizing abiotic parameters and passaging increased the fractional abundance of the target lineage, enabled definition of a nine-member core, and facilitated isolation of a *Terriglobus* sp. for downstream experimentation and morphological characterization by SEM. The same workflow revealed signals in other under-culture phyla, underscoring its generality for discovering substrates and partners for hard-to-culture taxa. Together, these results provide a tractable experimental framework that links genomes to processes in soils, supports targeted isolation and precision community editing [100], and supplies parameters and components for predictive models relevant to soil carbon cycling, plant productivity, and sustainable land management.

## Acknowledgments

We would like to thank Misun Kang and the staff at the University of California Berkeley Electron Microscope Laboratory for advice and assistance in electron microscopy sample preparation and imaging. We thank Matthew Incha and Mitchell Thompson for the development of the 96-well plate of phenolic compounds. This material by m-CAFEs Microbial Community Analysis & Functional Evaluation in Soils, a Science Focus Area led by Lawrence Berkeley National Laboratory, is based upon work supported by the U.S. Department of Energy, Office of Science, Office of Biological & Environmental Research under contract no.: DE-AC02-05CH11231.

## Conflict of interest

The authors declare that the research was conducted in the absence of any commercial or financial relationships that could be construed as a potential conflict of interest.

## Availability of data and materials

The raw sequence files for 16S rRNA gene amplicons and the *Terriglobus. sp.* DMC71 long-read sequencing on the NCBI Sequence Read Archive (SRA) under the accession SUB16156065 is underway and will be made available upon publication. The metagenomes from the longitudinal study will be uploaded to Figshare and made available upon publication. The *Terriglobus* sp. DMC71 strain submission to ATCC Type Strain Depository is underway. Scripts, ASV tables, and metadata for all microbiome related analysis are available at the github repository https://github.com/dawn-chiniquy/Chiniquy_2026.

## Notes

### Competing Interest Statement

The authors have declared no competing interest.

