## Supplemental Materials for "Metagenome-guided substrate selection enriches *Terriglobus*, reveals co-occurring taxa, and enables isolation of a novel species"

Short Title: CAZyme-guided *Terriglobus* Enrichment

Dawn Chiniquy[1]*, Spencer Diamond[2], Hans K. Carlson[1], Alexey Kazakov[1], Devin Coleman-Derr[3,4], Trent R. Northen[1,5], Jillian F. Banfield[2,6,7,8], Adam M. Deutschbauer[1,4]*

[1]Environmental Genomics and Systems Biology Division, Lawrence Berkeley National Laboratory, Berkeley CA; [2] Innovative Genomics Institute - University of California, Berkeley CA ;[3] Plant Gene Expression Center, USDA-ARS; [4]Department of Plant and Microbial Biology, University of California, Berkeley, CA; [5]Joint Genome Institute, Lawrence Berkeley National Laboratory, Berkeley CA; [6]Department of Earth and Planetary Sciences, University of California, Berkeley, CA; [7]Department of Environmental Science, Policy, and Management, University of California, Berkeley, CA; [8]Earth and Environmental Sciences, Lawrence Berkeley National Laboratory, Berkeley, CA

### Contents

This file contains Supplementary Methods, Supplementary Figures S1–S10 with legends, and Supplementary Tables S8 and S9. Supplementary Tables S1–S7, S10, and S11 are provided as separate files. Supplementary references are listed at the end of the Supplementary Methods.

### Supplementary Methods

*Identification of CAZy associated with rarely cultivated soil lineages from soil metagenome data*

Non-Archaeal MAGs with an estimated completeness ≥ 70% and contamination ≤ 5% using CheckM v1.2.1 [1] were retrieved (n = 558 bacterial MAGs) from the dataset of Diamond, et al. 2019 [2]. Proteins in MAGs were predicted with Prodigal v2.6.3 [3] and CAZyme were annotated using the dbCAN v6 HMM database [4]. Results were filtered to remove hits with an e-value $\geq 1 \times 10^{-14}$ and HMM coverage of ≤ 0.35. For CAZyme domains overlapping the same region of sequence, the domain with the lower e-value was selected. The total count of CAZyme per genome is available in Table S2.

Statistical analysis of CAZyme associations was performed in R v4.3.1. CAZy classes present in at least 2 MAGS were zero-imputed and normalized using centered log ratio (clr). Counts were normalized within each CAZy class to reflect the relative contribution of each MAG to the total pool of each CAZy class. PCA and UMAP were performed directly on the normalized matrix using the prcomp function in R and the umap function from the umap package in R, respectively. Permutational analysis of variance (PERMANOVA) was conducted using the adonis2 function of the vegan package in R. The association between the euclidean distance between genomes in the clr normalized CAZyme count matrix and if a genome was a member of one of four soil phyla recalcitrant to isolation (Acidobacteriota, Verrucomicrobiota, Chloroflexota, and Gemmatimonadota) was estimated using 9,999 permutations.

Association of CAZy classes with recalcitrant phyla was evaluated using generalized linear models in R of the form clr_normalized_cazy_abundance ~ recalcitrant, with recalcitrant being TRUE or FALSE for each of 558 MAGs. Significant differences between the set of recalcitrant and non-recalcitrant MAGs for each CAZy class were evaluated using the emmeans package [5] in R and p-values were adjusted using false discovery rate (FDR) with FDR ≤ 0.05 being significant.

*Field sample collection and processing*

Soil was collected in February 2020 from the south meadow field site at the Angelo Coast Range Reserve in Branscomb, California (39° 44′ 21.4″ N 123° 37′ 51.0″ W). Soil was collected using a manual coring device with a 1.5 in × 7 in cylindrical polycarbonate insert to remove 10 cm of soil at a time from an individual sampling bore. Soil from 0–10 cm of bore was placed in sterile bags on ice, then transported back to the lab and stored at 4 °C until further processing. The collected soil was passed through a #10 2 mm sieve to remove large particles and plant material.

*Selection of compounds for enrichment experiments*

In total 246 unique enzyme classes were analyzed for the following CAZyme superclasses (Aux Activity: AA, Carbohydrate Esterase: CE, Glycosyl Hydrolase: GH, and Polysaccharide Lyase: PL). Glycosyl transferases (GT) were omitted from the analysis as their activity is not specific to carbohydrate bond cleavage.

*Enrichment of microbial taxa with carbon compounds*

For the targeted enrichment experiments, each compound was dissolved in water and filter-sterilized at 2X concentration. For all compounds, to normalize between monosaccharides and polysaccharides, we used a set weight (0.5 g/40 ml for 2X solution) rather than molar weight for compounds unless they were difficult to dissolve, then they were dissolved at the highest concentration that would dissolve completely in water, including: wheat (1% or 1g/100 ml heated to 100 °C), galactomannan (25 mg/40 ml heated to 100 °C for 30 min), and glucuronoxylan (50 mg/40 ml heated to 100 °C for 30 min). The Freshwater Universal Media (FUM) (Table S10), minimal media containing buffer, salt, minerals, and vitamins was prepared at 4X, adjusting the pH to 5.5 and filter sterilizing. For the untargeted enrichment experiments, the carbon plate was also dissolved in filter-sterilized water, and the phenolic compound plate was dissolved in DMSO or water depending on the compound and as previously described [6–8] (**Table S11**). An 100X amino acid mixture was prepared to further support the growth of microbes on single carbon sources.

Field soil extractions used for inoculating enrichment cultures were completed as described previously [9,10], in brief, 4 g of field soil and 40 ml of sterile milliQ water were placed in a glass beaker with a stir bar at room temperature and mixed for 15 min and filtered through a sterile coffee filter to remove large particles. This soil extract was serially diluted 1:10 twice, five milliliters were transferred to a 50 ml tube with 45 ml of sterile milliQ water, and the tube was inverted to mix. Initial enrichment cultures were established in 2 ml 96-well deep well blocks and covered with an AeraSeal gas permeable film (Millipore Sigma), with a 400 µl working volume. Each well the culture was: 40 ul soil extract, 100 µl 4X FUM media, cycloheximide 100 µg/ml final concentration, 4 µl 100X amino acid solution, 200 µl 2X carbon compound solution, and brought up to 400 µl with sterile water. For salt-stressed enrichment cultures, 37.4 g of NaCl was added to 200 ml 4X media, mixed to dissolve, then filter-sterilized. Once inoculated, cultures were grown in the dark at 30°C with 150 rpm shaking for 10 d. For time series experiments, each transfer to new media was a 1:10 subculture, transferring 40 µl of enrichment culture to 360 µl of new growth media. Samples that were collected for DNA isolation were centrifuged at full speed for 10 min, and pellets were stored at -20°C until DNA extraction. With each transfer and sample collection for DNA isolation, 20% glycerol stocks were made of the cultures to be able to go back to individual samples that had the highest abundance of difficult to culture microbes once profiled by 16S rRNA sequencing.

*DNA extraction, Amplification, and Illumina Sequencing of 16S Sequences*

Genomic DNA was extracted from samples using the 96-well Qiagen DNeasy Blood & Tissue kit (catalog no. 69516), following the manufacturer's protocol for Gram-positive bacteria. The V4/V5 region of the bacterial 16S rRNA gene was PCR amplified from extracted DNA using the 515F/926R primers with in-line dual Illumina indexes [11]. The resulting libraries were quantified using the Qubit dsDNA BR Assay kit (Invitrogen, catalog no. Q32853), pooled, cleaned up using the Zymo Research Clean and Concentrator kit (Zymo Research, catalog no. D4014), and sequenced on an Illumina MiSeq (PE 2x300). For 16S read processing, paired-end reads were merged using PEAR [12], demultiplexing using inline indexes, and reads with more than one expected error was detected and discarded, as well as rare sequences (fewer than 4 reads in any sample) using USEARCH [13]. Chimera removal and error correction were performed with UNOISE3 [14]. Taxonomy was assigned using SINTAX [15] with the RDP training set v18 [16].

*Assembly, binning, dereplication, and genome quality*

Raw metagenomes from 89 samples were assembled with IDBA-UD [17] and SPAdes [18]. Contigs were binned using four independent binners, and bins were consolidated with DASTool [19]. For each bin, taxonomy was assigned with GTDB-Tk [20], and completeness and contamination were estimated with CheckM [1]. Species-level dereplication used dRep [21] at ANI ≥95% to select a single species-representative bin per cluster, maximizing completeness and contiguity while minimizing contamination. We predicted coding sequences with Prodigal [3], rRNA and tRNA genes with standard workflows, and recorded codon usage with Codetta [22]. The final set comprised 106 species-representative genomes with high quality (mean completeness 95.9%, mean contamination 1.15%; **Table S7**).

*Read mapping and abundance profiling*

Quality-controlled reads were competitively mapped to the 106-genome representative set to obtain per-species coverage and relative abundance per sample. Mean mapping rates were ~80% across the dataset, consistent with expectations for reduced-complexity enrichment communities. To compare enrichments and derived stocks, we computed alpha diversity (observed species) and constructed ordinations on CLR-transformed relative abundance matrices.

*Core microbiome definition and network inference*

We defined prevalence as the fraction of enrichment samples (n=71) in which a species was detected. The core microbiome was the set of species present in ≥95% of enrichments. We also reported species present in 100% of enrichments and compared core membership between biological replicates of enrichment communities over time. We filtered to species present in ≥10 enrichments (n=71) to reduce sparsity. We inferred two complementary networks: SparCC correlations from relative abundance data with permutation to assess edge significance [23] and SPIEC-EASI conditional-dependence graph [24] using neighborhood selection with StARS for model selection [25]. We retained positive edges and defined a consensus network by requiring presence in both SparCC and SPIEC-EASI [24,25]. We summarized node-level statistics (degree, betweenness, eigenvector centrality) and global properties using igraph [26]. To identify community structure, we applied the weighted Leiden algorithm [27] with modularity optimization across resolution values ranging from 0.1 to 1.5 (step = 0.05), selecting the resolution that maximized modularity (resolution = 0.75, modularity = 0.67). To assess clustering stability, we performed 100 independent Leiden iterations at the optimal resolution using different random seeds; 79% of runs recovered seven communities with consistent modularity scores (mean = 0.67 ± 0.004). We highlighted direct neighbors of Acidobacteriota and compared their taxonomic composition to the overall species set.

*CAZyme annotation of enrichment MAGs*

To assess hemicellulose-degrading capacity among direct network neighbors and core microbiome members, we predicted proteins with Prodigal v2.6.3 and searched against Pfam HMMs for GH43 (PF04616) and GH62 (PF03664) using HMMER v3.4 with an E-value threshold of $1 \times 10^{-5}$.

*Isolation of Terriglobus sp. from enrichment cultures*

Enrichment cultures were re-grown from glycerol stocks of the communities stored in a -80 °C freezer by inoculating 3 mls of FUM minimal media pH 5.5 with glucuronoxylan (50 mg/40 ml) with growth for 10 d, serially diluted to 10-4 and spread on agar plates composed their respective growth media. Individual colonies were collected and re-streaked on the same media. Individual colonies from previously re-streaked colonies were grown in 0.1X R2A for 5 d, then pelleted, and DNA was isolated for 16S profiling. Mixed cultures identified through sequencing would be re-streaked on minimal media again before being grown in 0.1X R2A for 5 d, pelleted, and DNA was isolated for another round of 16S profiling. Once cultures were considered pure, isolates were grown in 3 ml of 0.1X R2A for 5 d (or until cloudy) for DNA isolation and metagenome sequencing. Isolation of the *Terriglobus* sp. DMC71 required over 6 rounds of re-streaking to get a pure culture.

*Long-read genome sequencing and classification of Terriglobus sp. DMC71*

Hybrid long-read sequencing (Oxford Nanopore; Plasmidsaurus) produced a closed, circular chromosome of 6.15 Mb with 57.9% GC content, 100% completeness, and 0% contamination (CheckM). The *Terriglobus* sp. DMC71 genome was classified with GTDB-Tk v 2.4.0 against the Genome Taxonomy Database (GTDB) release 220 [28,29].

*SEM imaging of Terriglobus sp. DMC71*

Scanning electron microscopy preparations (Electron Microscopy Laboratory, University of California, Berkeley) were: *Terriglobus* sp. DMC71 isolate was grown in 0.1X R2A for 7 d (shaking at 150 rpm at 30°C, then gently pelleted, and resuspended in a fixative solution (2% glutaraldehyde in 0.1 M sodium cacodylate buffer, pH 7.2) and was placed at 4°C rotating for 3 hr then stored at 4°C in dark (standing) overnight. Prior to imaging, the sample was washed 3x in 0.1 M sodium cacodylate buffer, pH 7.2, postfixed 3x in 1% osmium tetroxide in 0.1 M sodium cacodylate buffer, pH 7.2 for 1 h, and then washed 3x in 0.1 M sodium cacodylate buffer, pH 7.2. Samples were dehydrated in a graded ethanol series up to 100%. Samples were then transferred onto 0.1% (w/v) poly-L-lysine–coated coverslips and subjected to critical point drying (Tousimis, Mayland, USA). The dried coverslips were mounted onto aluminum stubs using conductive carbon tape and sputter-coated with a gold/palladium alloy (Safematic GmbH, Switzerland). SEM imaging was performed using a Zeiss Crossbeam 550 (Carl Zeiss Microsystems GmbH, Oberkochen, Germany).

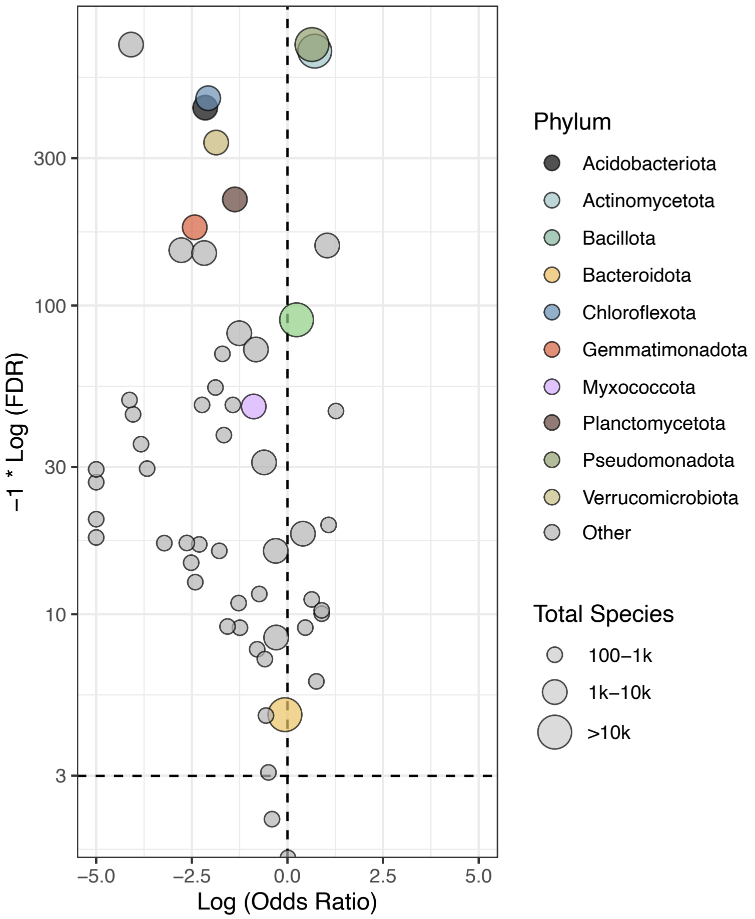


**Figure S1. Prevalence of Latinized species names across globally abundant soil phyla.** Volcano plot showing the prevalence of Latinized species names in the Genome Taxonomy Database (GTDB-R226) across globally abundant soil phyla, used here as a proxy for species-level isolation. The x-axis shows log odds ratio (LOR), where negative values indicate under-representation in named isolates and positive values indicate over-representation. The y-axis shows –log10(FDR), and point size reflects the total number of GTDB species assigned to each phylum. Dashed lines mark LOR = 0 and FDR = 0.001. Acidobacteriota, Verrucomicrobiota, Chloroflexota, Gemmatimonadota, and Planctomycetota were significantly under-represented among named isolates (LOR < 0, FDR ≤ 0.001) and were treated as recalcitrant phyla throughout this study.

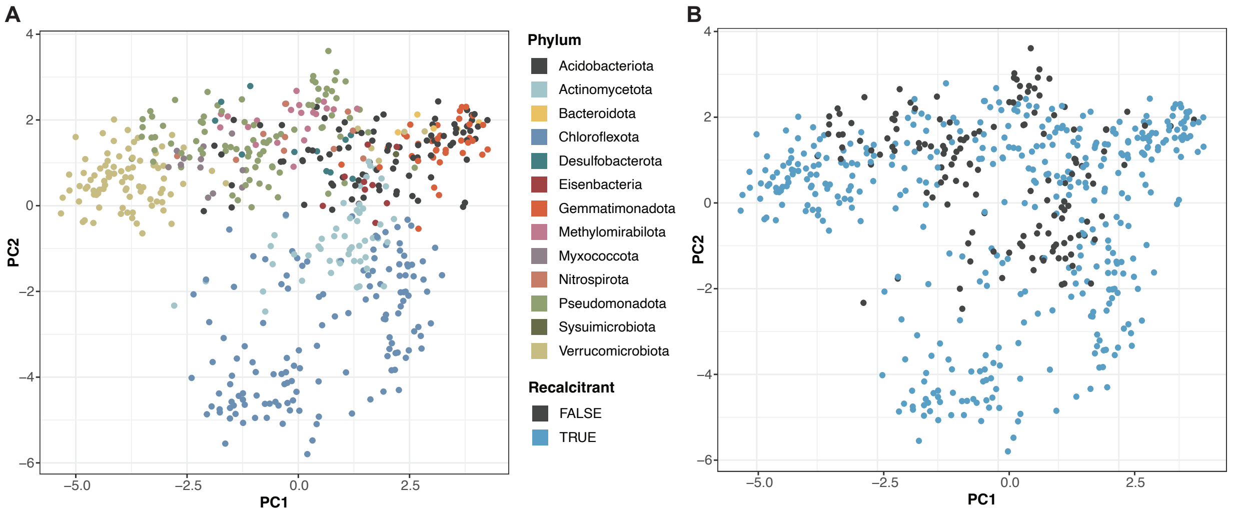


**Figure S2. Principal component analysis of CAZyme repertoires across soil MAGs.** Principal component analysis (PCA) of centered log-ratio (CLR)-normalized CAZyme counts across 558 high-quality bacterial MAGs (>70% completeness, <5% contamination) recovered from Angelo Coast Range Reserve grassland soil metagenomes. Each point represents one MAG. **(A)** MAGs colored by GTDB phylum. **(B)** Same ordination colored by whether the phylum was over-represented or under-represented in named isolate collections (recalcitrant: False/True). The ordination shows broad phylum-level separation in a horseshoe pattern, indicative of a non-linear relationship.

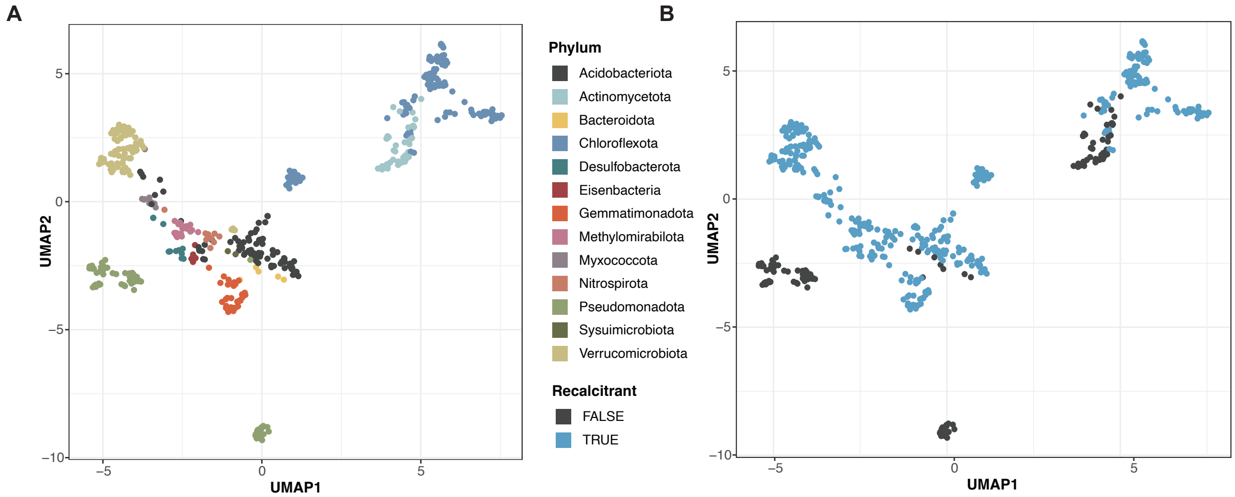


**Figure S3. CAZyme repertoires separate soil MAGs by phylum and by representation in culture collections.** Centered log-ratio (CLR)-normalized CAZyme counts from 558 high-quality bacterial MAGs (>70% completeness, <5% contamination) recovered from Angelo Coast Range Reserve grassland soil metagenomes were visualized with UMAP. Each point represents one MAG. **(A)** Points are colored by GTDB phylum. Panel A reproduces the UMAP shown in Figure 2A to facilitate comparison with panel B, which colors the same ordination by recalcitrant status. **(B)** The same ordination is colored by whether the phylum was over-represented or under-represented in named isolate collections (recalcitrant: False/True). Together, the two panels show that CAZyme composition captures both taxonomic structure and the contrast between recalcitrant and non-recalcitrant lineages.

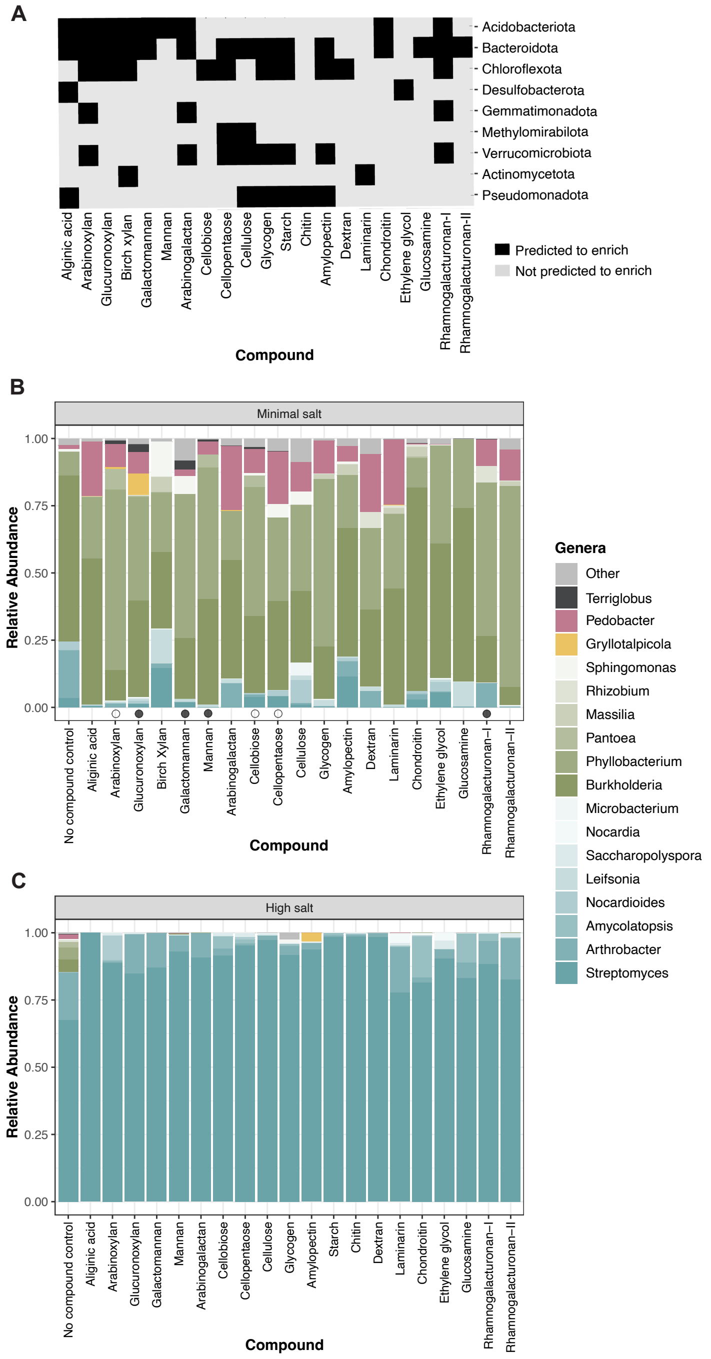


**Figure S4. Predicted substrates and salt-dependent enrichment outcomes in the targeted screen. (A)** Summary of Table S6 showing predicted substrate enrichments by phylum based on CAZyme analysis and commercial substrate availability. **(B)** Genus-level relative abundance profiles for enrichments grown under minimal-salt conditions. (**C**) Genus-level relative abundance profiles for enrichments grown under high-salt conditions. Open and filled circles adjacent to compound names indicate conditions with one replicate or multiple replicates, respectively, containing detectable Acidobacteriota reads. High-salt conditions favored Actinomycetota, whereas low levels of Acidobacteriota were detected only in a subset of compounds under minimal-salt conditions.

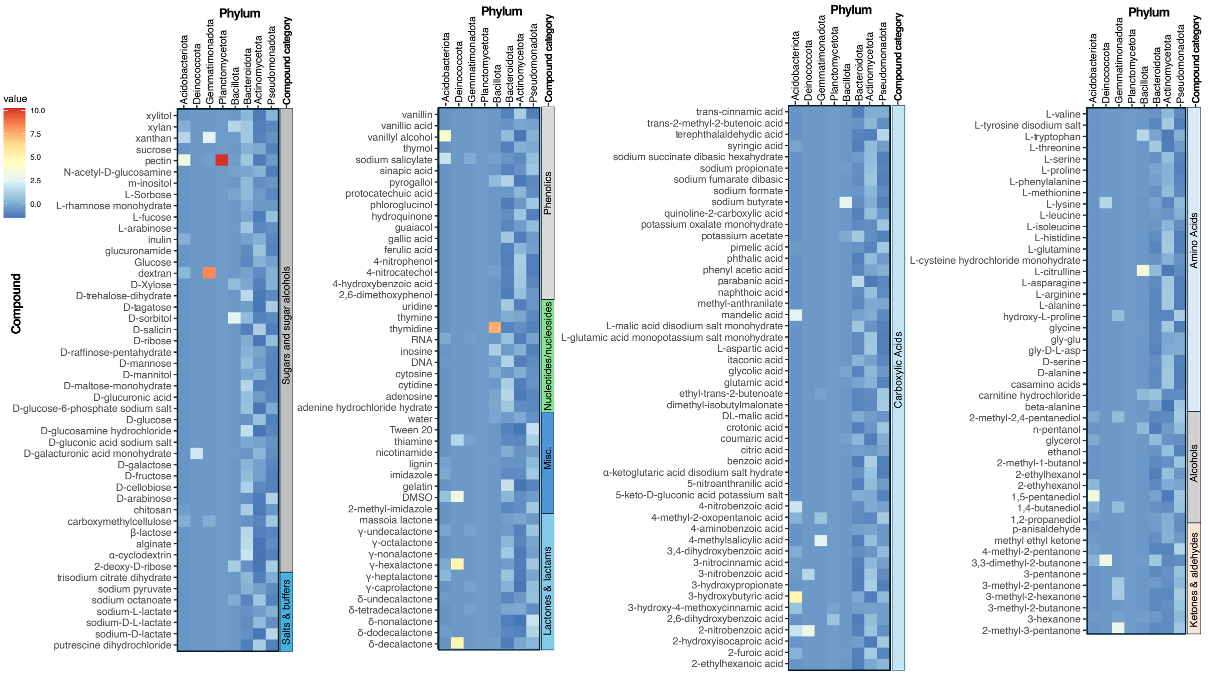


**Figure S5. Untargeted 192-compound enrichment screen.** Heat map showing 16S rRNA amplicon sequencing results from soil enrichments grown on 192 compounds distributed across a carbon plate and a phenolic compound plate, each run on four replicate plates. Each tile represents the mean relative abundance across four biological replicates for a given compound. The screen was performed to benchmark the CAZyme-guided targeted approach against a broader untargeted substrate survey.

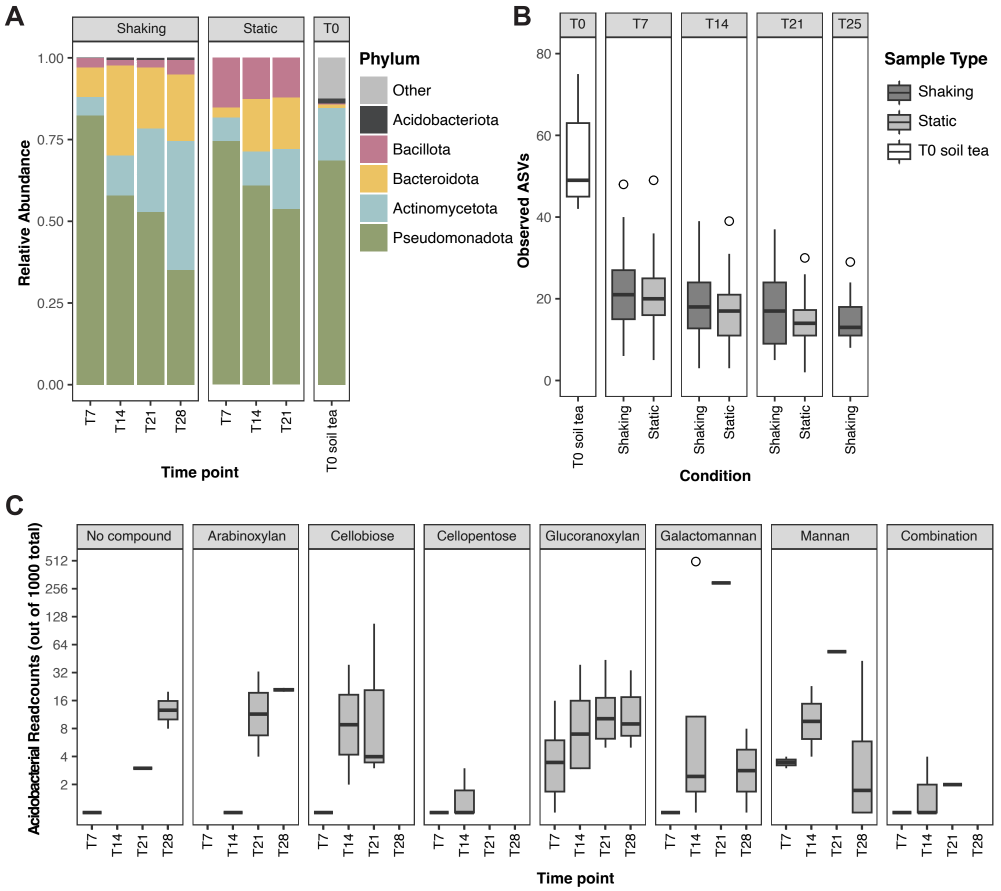


**Figure S6. Retesting and optimization of Acidobacteriota-enriching conditions. (A)** Phylum-level relative abundance profiles from 16S rRNA amplicon sequencing comparing shaking and static growth across the optimization time course. Acidobacteriota were detected only in shaken cultures. T0 soil tea is shown for reference. **(B)** Observed ASVs across the same shaking and static treatments. **(C)** Acidobacteriota read counts over time (log scale) for the six individual substrates retested for Acidobacteriota enrichment and for the admixture of all six substrates. Acidobacteriota enrichment was generally favored at days 14 and 21, and the substrate admixture did not improve enrichment relative to the best-performing individual substrates.

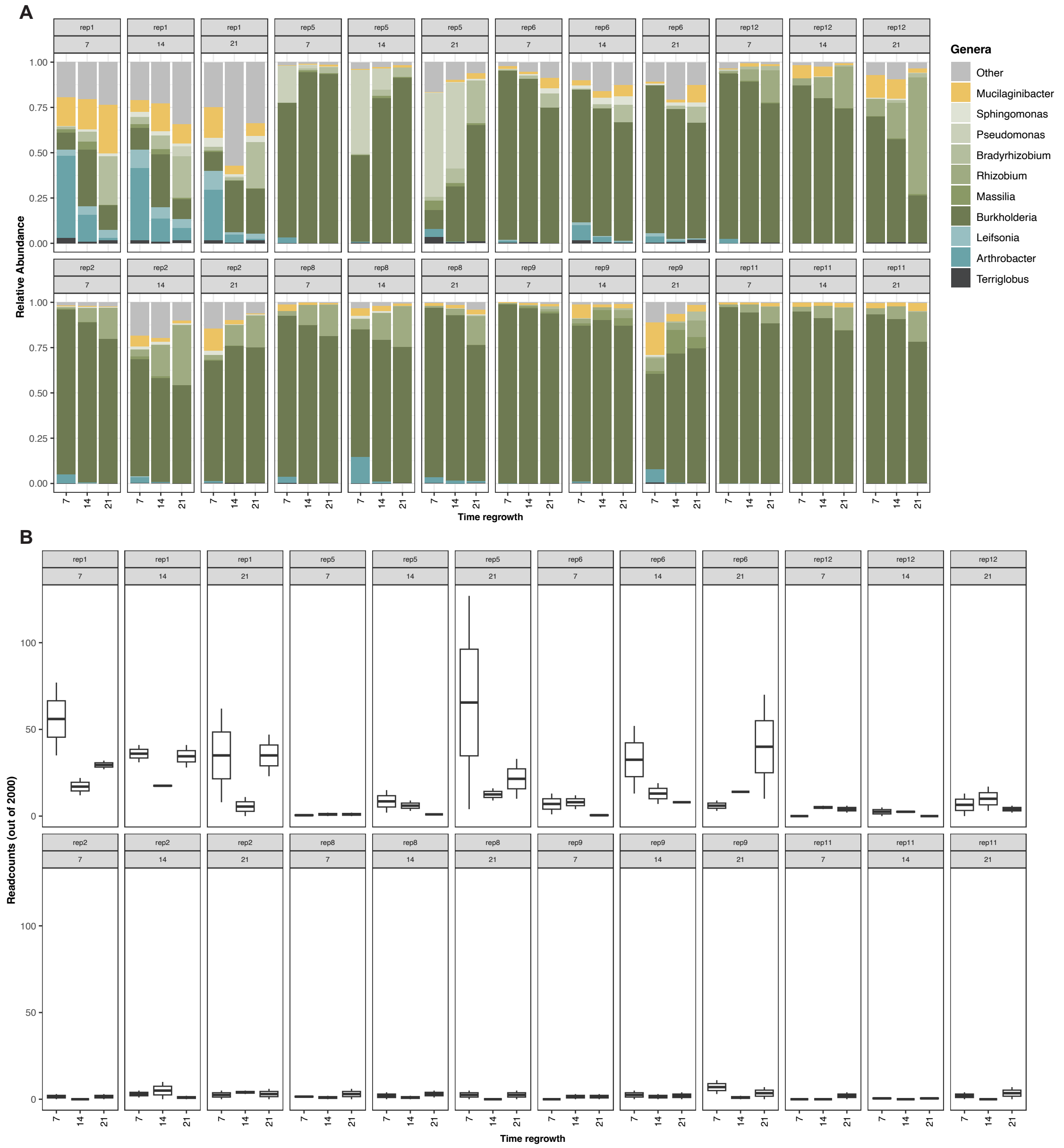


**Figure S7. First-stage passaging of glucuronoxylan-derived communities prior to longitudinal metagenomics.** To generate compositionally diverse inocula for downstream shotgun metagenomics, eight glucuronoxylan-derived enrichment communities were selected from Figure 5A, including four with high Acidobacteriota abundance and four with lower abundance. Communities were passaged and sampled at days 7, 14, and 21, and glycerol stocks from each timepoint were used as inocula for second-stage passaging (Figure 6). **(A)** Relative abundance profiles over time based on 16S rRNA amplicon sequencing. **(B)** Acidobacteriota read counts (out of 2,000 total reads) for high-abundance communities (top) and low-abundance communities (bottom).

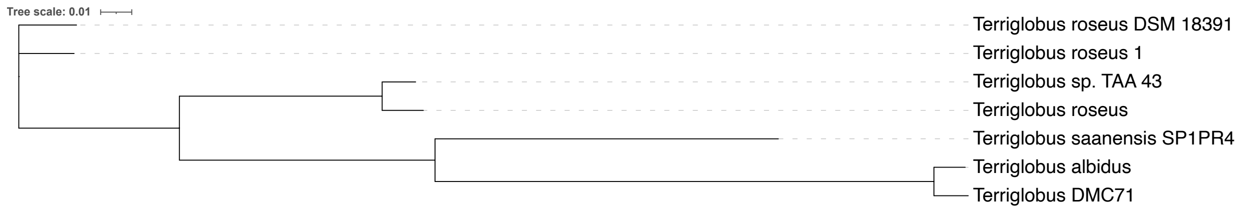


**Figure S8. Phylogenetic placement of *Terriglobus sp.* DMC71.** Phylogenetic tree of Terriglobus sp. DMC71 inferred with FastTree from a concatenated alignment of 120 bacterial marker proteins generated by GTDB-Tk. *Terriglobus sp.* DMC71 clusters closest to *Terriglobus albidus*, its nearest named relative, consistent with the ANI-based comparison reported in the main text. Support values were 1.0 for all internal nodes.

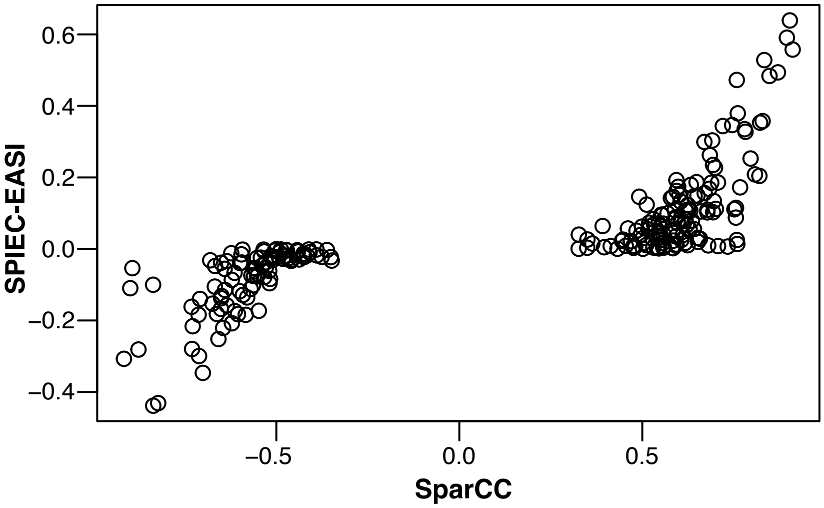


**Figure S9. Agreement between network inference methods.** Correlation of edge weights between SparCC and SPIEC-EASI (neighborhood selection) co-occurrence networks. Only edges detected by both methods are shown (n = 232). The strong Spearman correlation ($\rho = 0.916$, $p < 2.2 \times 10^{-16}$) indicates high agreement between methods. Edges with conflicting signs were removed prior to downstream analysis.

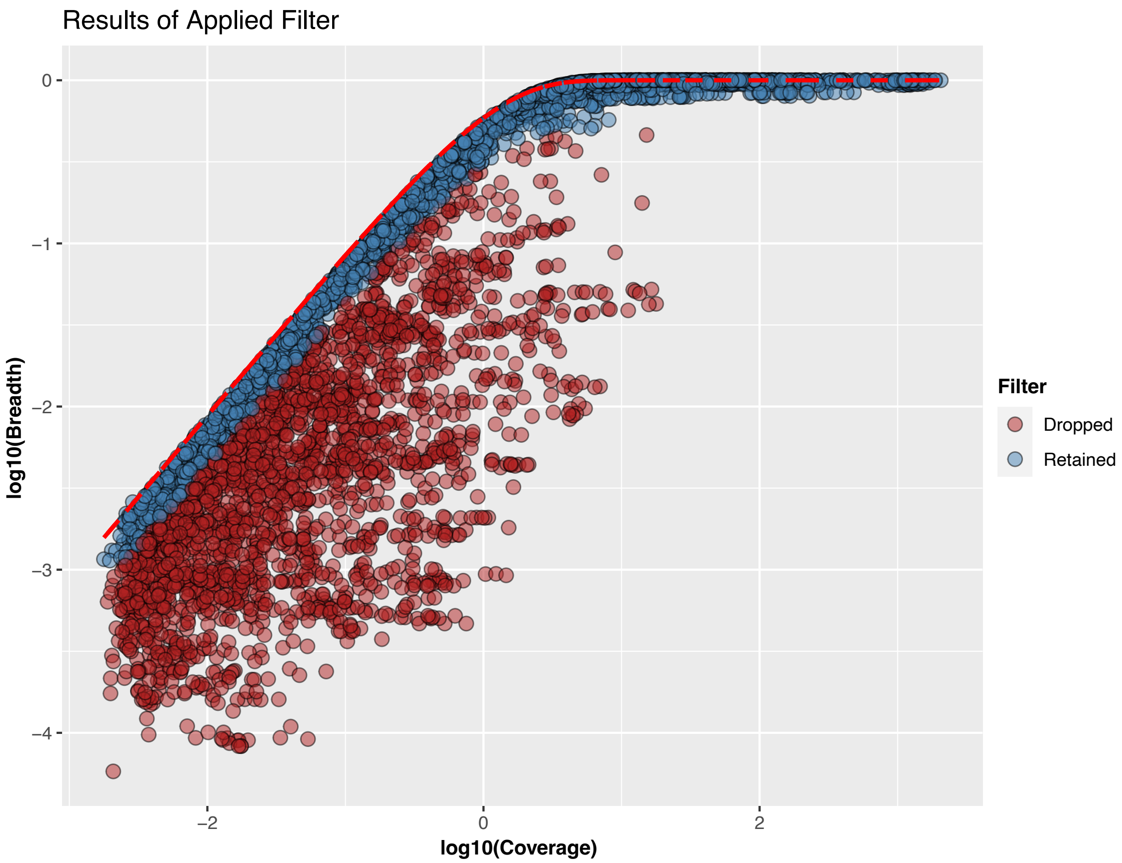


**Figure S10. Coverage and breadth metrics for MAG filtering.** Relationship between genome coverage and breadth across enrichment samples for the 106 MAGs included in downstream network analysis. MAGs meeting filtering thresholds of breadth > 0.5 and coverage > 1× were retained for co-occurrence analysis.

**Table S8. Core microbiome members across enrichment samples and their GH43 family content.** Nine species met the core-microbiome threshold (detected in ≥95% of n = 71 enrichment samples) and are listed with their GTDB-R226. Detected indicates the percentage of enrichment samples in which each genome was recovered above breadth and coverage thresholds (breadth > 0.5; coverage > 1×; see Supplementary Methods).

| Genome | Detected | Phylum | Class | Order | Family |
|---|---|---|---|---|---|
| Acido – Core 1 | 100 | p__Acidobacteriota | c__Acidobacteriae | o__Acidobacteriales | f__Acidobacteriaceae |
| Core 2 | 100 | p__Proteobacteria | c__Alphaproteobacteria | o__Caulobacterales | f__Caulobacteraceae |
| Core 3 | 100 | p__Proteobacteria | c__Alphaproteobacteria | o__Rhizobiales | f__Xanthobacteraceae |
| Core 4 | 100 | p__Proteobacteria | c__Alphaproteobacteria | o__Rhizobiales | f__Xanthobacteraceae |
| Core 5 | 100 | p__Proteobacteria | c__Alphaproteobacteria | o__Rhizobiales | f__Xanthobacteraceae |
| Core 6 | 100 | p__Proteobacteria | c__Alphaproteobacteria | o__Rhizobiales | f__Xanthobacteraceae |
| Core 7 | 100 | p__Proteobacteria | c__Alphaproteobacteria | o__Rhizobiales | f__Xanthobacteraceae |
| Core 8 | 98.5915493 | p__Proteobacteria | c__Alphaproteobacteria | o__Rhizobiales | f__Rhizobiaceae |
| Core 9 | 97.1830986 | p__Proteobacteria | c__Alphaproteobacteria | o__Rhizobiales | f__Rhizobiaceae |

**Table S9. GH43-encoding genomes among direct network neighbors of *Terriglobus*.** Direct co-occurrence partners of the *Terriglobus_A* core member with positive, sign-consistent edges in the SparCC/SPIEC-EASI consensus network (see Supplementary Methods) that encode glycoside hydrolase family 43 (GH43) enzymes, listed with their full enrichment-MAG identifier, GTDB-R226 taxonomy, core-microbiome status (TRUE/FALSE; see Table S8), and total GH43 count. The non-core *Niastella* MAG (Bacteroidota; family Chitinophagaceae) encodes 13 GH43 homologs, which is more than any core member, including *Terriglobus*, nominating it as a second major hemicellulose degrader co-occurring with *Terriglobus* across the longitudinal series. Genome identifiers are abbreviated from the full enrichment-MAG IDs (prefix "AngeloEnr2023_" omitted); full identifiers are provided in the standalone supplementary data files.

| **Genome** | **Taxonomy (Phylum; Class; Order; Family)** | **Genus** | **Core** | **GH43** |
|---|---|---|---|---|
| xylan_42_C05_OT19_TP14_spades_concoct_25_sub | Bacteroidota; Bacteroidia; Chitinophagales; Chitinophagaceae | *Niastella* | FALSE | 13 |
| xylan_02_C01_OT13_TP07_spades_metabat.16 | Proteobacteria; Alphaproteobacteria; Caulobacterales; Caulobacteraceae | BOG-938 | TRUE | 1 |
| lbgedit_85_MCD_GL30_TRA_spades_concoct_13 | Proteobacteria; Alphaproteobacteria; Rhizobiales; Xanthobacteraceae | *Bradyrhizobium* | TRUE | 0 |
| lbgedit_82_MCC_GL30_TRA_spades_metabat.6 | Proteobacteria; Alphaproteobacteria; Rhizobiales; Xanthobacteraceae | *Bradyrhizobium* | TRUE | 0 |
